# Serial intravital microscopy reveals temporal dynamics of autoreactive germinal centers in the spleen

**DOI:** 10.1101/2025.01.06.631523

**Authors:** Layla Pohl, Thomas R. Wittenborn, Ali Shahrokhtash, Kristian S. Kastberg, Cecilia Fahlquist-Hagert, Lisbeth Jensen, Donato Sardella, Alain Pulfer, Duncan Sutherland, Santiago F. Gonzalez, Ina Maria Schiessl, Søren E. Degn

## Abstract

The spleen plays a key role in clearing blood-borne infections and is involved in autoimmune and hematological disorders. It undergoes extensive remodeling during inflammation and immune reactions, but its localization in the peritoneal cavity has hampered studies of these dynamic changes. Here, we establish and validate a protocol for serial 2-photon microscopy of the murine spleen to capture dynamic processes in the living animal. As a proof-of-principle, we elucidate the expansion and contraction of autoreactive germinal centers (GCs) induced by epicutaneous application of the small-molecule TLR7 agonist resiquimod (R848). Leveraging a biocompatible abdominal imaging window, intravital labeling techniques, and fluorescent reporters, we follow GCs up to 180 µm below the capsule for more than 2 weeks by tracking follicular dendritic cell (FDC) networks. This was accomplished without appreciable perturbation of normal physiology, paving the way for a deeper understanding of the biology of the spleen and its associated disease states.

**Highlight:** An abdominal imaging window allowing the study of dynamic processes in the spleen of live mice over the course of several weeks.

## Introduction

The spleen is the largest secondary lymphoid organ. It harbors two morphologically and functionally distinct compartments, the red pulp and the white pulp, which are separated by the marginal zone. In the white pulp, B cells are organized in follicles surrounding the periarteriolar lymphoid sheath (PALS), which contains dendritic cells and T cells. The red pulp and the marginal zone contain macrophages which filter out damaged and senescent erythrocytes, as well as foreign material, whereas the white pulp orchestrates adaptive immune responses to the captured antigens (*1*). Hence the spleen plays a key role in clearing blood-borne infections and is involved in numerous systemic conditions including autoimmune diseases such as systemic lupus erythematosus (SLE), and hematological disorders (*2*).

The spleen undergoes extensive remodeling during inflammation and immune reactions, but due to its localization in the peritoneal cavity, it has traditionally been challenging to study these dynamic changes. Prior work using intravital microscopy in mice elucidated the transport of immune complexes across the marginal zone to the underlying follicles (*3*), a key step in antigen presentation, and the influx of T cells along perivascular-T (PT)-tracks during an active immune response (*4*). However, these studies relied on open surgery to expose the spleen and were limited to timespans of hours, an approach which did not permit longitudinal tracking of dynamic immune processes occurring over days to weeks.

Longitudinal studies of immune processes necessitate the avoidance of exogenous proinflammatory contributors, immune provocations, and disturbances of natural lymphatic and blood flows bringing in antigens and immune cells. This has been achieved with imaging windows for superficial cutaneous lymph nodes to study cancer metastases (*5–7*), germinal center (GC) B-cell responses (*8*), and follicular T-cell subsets (*9*). A similar setup with implantation of an imaging window in the abdominal wall has also been developed for studying physiological processes in internal organs (*10, 11*). This abdominal imaging window (AIW) has been leveraged to interrogate the dynamic function of organs in the abdominal cavity such as the kidney (*12, 13*), the pancreas (*14*), and the liver (*15, 16*). The spleen, however, is a particularly challenging organ because of its high blood-flow, thick capsule, high density of active and autofluorescent immune cell populations (*17*), and its ability to dynamically expand and contract over a very large size range.

Here, we describe the development of a robust protocol for functional studies in the spleen over several weeks using an AIW. We leverage an anti-fouling approach to render the AIW surface biocompatible and demonstrate that the presence of the AIW does not perturb the normal immune homeostasis. We demonstrate its utility by tracking follicular dendritic cell (FDC) network expansion and contraction in the context of an autoimmune response, utilizing a pharmacological model of SLE based on epicutaneous application of the small-molecule Toll-like receptor (TLR)-7 agonist resiquimod (R848) (*18*).

TLR7 is a well-known driver of autoimmunity in mice (*19–22*), and TLR7 gain-of-function mutation has been found to cause SLE in humans (*23*). SLE is characterized by the production of affinity-matured autoantibodies, which can originate from both extrafollicular and GC-driven B-cell responses (*24, 25*). While extrafollicular responses are sufficient for the early hallmarks of lupus (*26*), GC responses are thought critical to epitope spreading and progression of disease (*27*). Both patients with, and mouse models of, SLE present with spontaneous GCs in the spleen (*28–30*), and autoreactive GC responses are known to depend upon B-cell intrinsic TLR7 signaling (*31–33*). TLR7 sensing by FDCs also promote GC responses by provision of type I interferon (*34*), which drives autoreactive B-cell activation (*35, 36*).

In GCs, B-cell centroblasts proliferate in the dark zone and undergo somatic hypermutation of their immunoglobulin genes to introduce subtle variations in antigen recognition (*37*). They subsequently migrate into the light zone to test their affinity for antigen retained in intact form by FDCs via complement receptor 1 (CR1, CD35) and Fc-gamma receptor IIb (FcγRIIb) (*38*). B-cell centrocytes that can competitively take up antigen and present antigen-derived peptides to T-follicular helper cells (T_FH_) are selected for a renewed round of expansion and hypermutation or for plasma cell or memory B-cell differentiation. B cells that are unable to bind antigen undergo apoptosis and are phagocytosed by highly active tingible-body macrophages (TBMs).

Leveraging our AIW, we show that the dynamic expansion and contraction of GCs occurring in connection with initiation and resolution of an autoimmune inflammatory response can be followed by tracking the FDC network. We further demonstrate that we can achieve cellular resolution intravitally using the longitudinal approach. Our methodology opens the door to future studies of dynamic spleen functions in health and disease.

## Results

We developed a novel surgical protocol adapted from the previously described AIW implantation over the left kidney (*39*) to allow imaging of the spleen. To this end, we employed a modified titanium window (*40*) that was retailored from the original AIW model (*41*) to specifically enable and simplify serial intravital imaging of abdominal organs using an upright 2-photon microscope.

Encapsulation of organs by connective tissue over time is a known problem when implanting AIWs, pushing the organs deeper into non-imageable depths. (*42*). This problem can be explained by a foreign-body response including both acute and chronic inflammation and the development of granulation tissue (*43*). Because the spleen is a highly active immune organ, we anticipated the necessity of passivating the AIW surface to prevent such formation of granulation tissue impeding imaging and impacting the systemic immune response.

### High-temperature PMOXA coating reduces immunogenicity of the AIW

We aimed to reduce the foreign-body response by glass coverslip passivation to establish longitudinal intravital imaging conditions that perturb the immune system minimally. We chose a coating consisting of oxidation-stable antifouling brushes of 4.25 kDa Poly-2-methyl-2-oxazoline (PMOXA) (*44, 45*), with predefined spacing, grafted on a polyacrylamide (PAcrAm) backbone. High-density PMOXA brushes were grafted to the coverslip surface using high-affinity electrostatic amine linker and covalent silane linkers designed on PAcrAm backbone for long-term stability (*46*). To test different passivation techniques, we evaluated murine macrophage and fibroblast adherence on the glass coverslip over 2 weeks; macrophages, as they are key early responders, and fibroblasts as they contribute later in foreign-body responses with extracellular matrix deposition and tissue contraction during the encapsulation process (*43, 47*). Accordingly, we evaluated cellular overgrowth of murine 3T3 fibroblasts and RAW 264.7 macrophages on non-coated glass and two different passivation processes, room temperature PMOXA and high-temperature PMOXA coating. Culturing these cells for 2 weeks on the passivated substrates demonstrated that the high-temperature PMOXA passivation protocol resulted in fewer adherent cells for both cell lines (Fig. 1A and B). Additionally, cells exhibited reduced spreading on the passivated surfaces, which provided a clearer field of view (Fig. 1C). Consequently, the high-temperature PMOXA passivation coating was selected as the standard coating protocol.

**Fig. 1.**
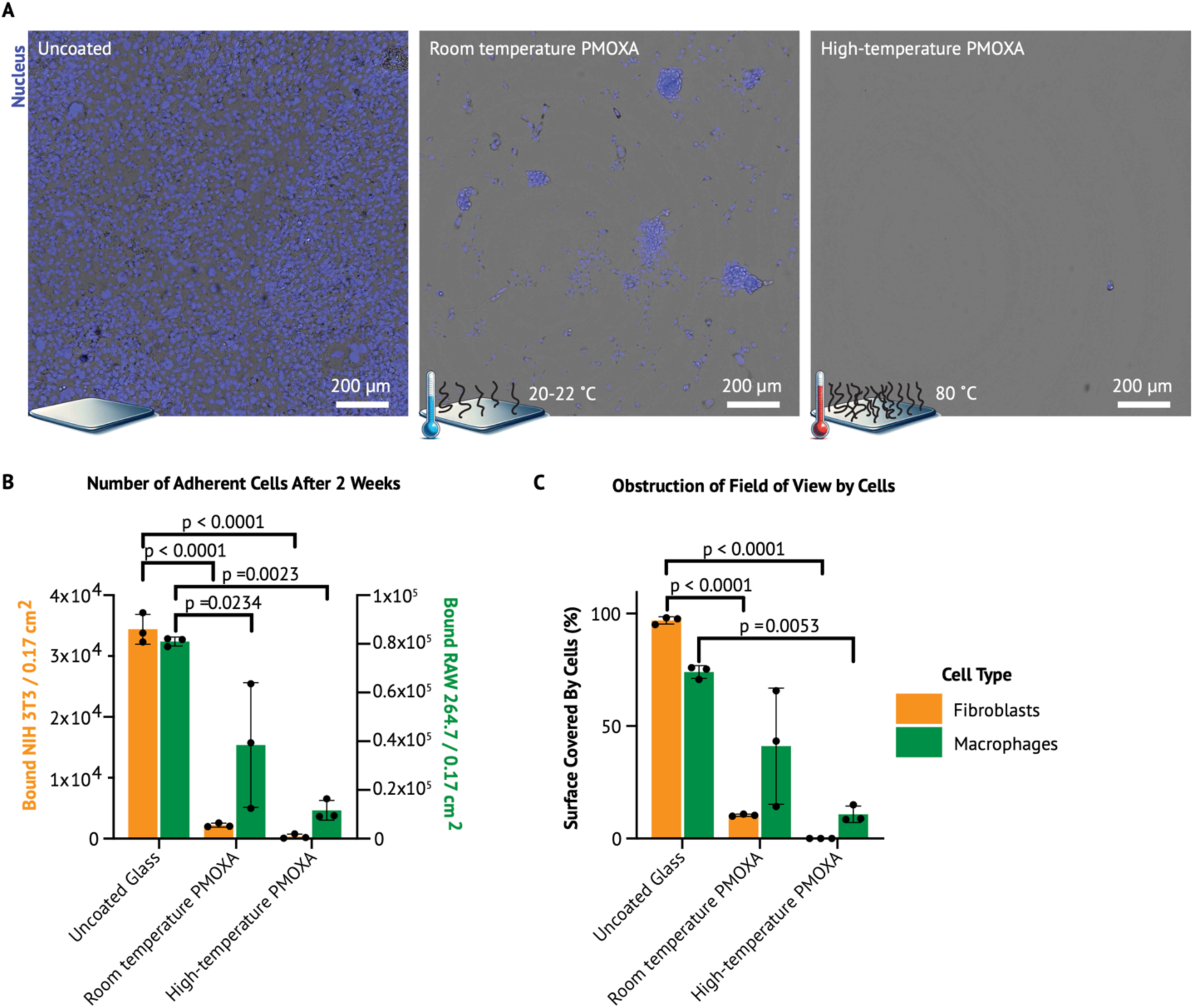
*In vitro* comparison of different PMOXA passivating coatings after 2 weeks. **(A)** Representative brightfield and nucleus-stained images of fibroblasts adhering and spreading on uncoated and passivated substrates after 2 weeks in serum-containing media. The uncoated substrates became overgrown within a few days, whereas room temperature PMOXA passivation significantly reduced non-specific cell adhesion, leading to the formation of 3D colonies on the few adherent cells. High-temperature PMOXA passivation almost entirely prevented fibroblast adhesion. **(B)** Quantification of the number of adherent cells for each passivation condition. High-temperature PMOXA passivation was significantly more effective in preventing non-specific adhesion of both fibroblasts and macrophages. **(C)** Quantification of surface area covered by cells on different substrates. High-temperature PMOXA passivation resulted in a clearer field of view due to a lower number of adhered cells and reduced cell spreading on the surface. Data are presented as means ± SD. P values were calculated using ordinary one-way ANOVA, comparing each condition to the uncoated control (n = 3 with 2 technical repeats).

Having established non-immunogenic conditions *in vitro*, we sought to implement the AIW for spleen imaging *in vivo*. The spleen is a key player in systemic autoimmune diseases including SLE. Both autoimmune patients and mouse models of SLE display prevalent GC responses in the spleen and robust splenomegaly. We surmised that the dramatic remodeling of the spleen occurring in connection with an autoimmune response would at the same time present a good proof-of-principle and a challenging test case due to the expansion and contraction of the tissue. To be able to test, down the road, the feasibility of longitudinal imaging at cellular resolution, we chose FoxP3-DTR-GFP reporters, marking T regulatory (T_REG_) cells in general and the GC-relevant T-follicular regulatory (T_FR_) cell subset in particular.

### An AIW allows tracking autoreactive GC dynamics in the spleen over 2 weeks

FoxP3-DTR-GFP mice were treated with epicutaneous application of the small-molecule TLR7 agonist resiquimod (R848) (*48*) to commence an SLE-like autoreactivity. The AIW was implanted over the spleen (Fig. 2A and B) and the initiation of individual autoreactive GCs was followed through the AIW from R848 treatment commencement (Day 1) and for 2 weeks (Day 5-20) (Fig. 2C and D). To similarly follow the contraction of the response associated with remission, another cohort was treated with R848 for 4 weeks, then treatment was stopped, and individual GCs were followed for 2 weeks (Day 29-44) (Fig. 2C and E).

**Fig. 2.**
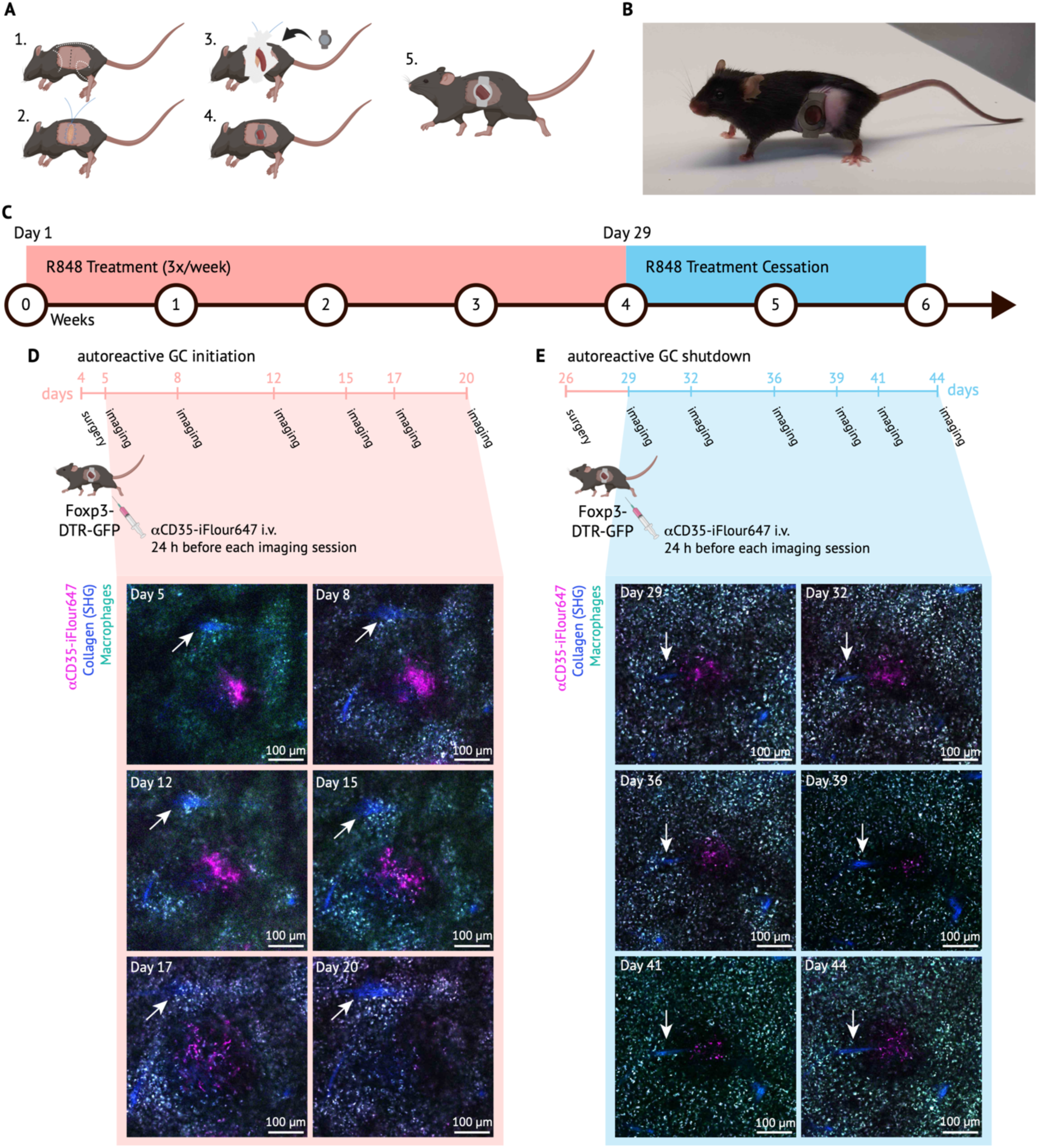
Serial intravital imaging of autoreactive germinal centers in the spleen. (**A**) Graphical overview of surgery steps: 1. AIW placement, 2. purse string suture, 3. spleen mobilization, 4. placement of AIW, 5. recovered mouse after surgery. (**B**) Recovered mouse 8 days after AIW implantation. (**C**) Global treatment timeline: Foxp3-DTR-GFP mice were treated with R848 three times per week for four weeks to induce an autoimmune response. Then treatment was stopped, and the contraction of the autoimmune response was followed for two weeks. (**D**) To investigate the emergence of autoreactive GCs, mice were implanted with an AIW shortly after R848 treatment initiation and individual GCs were followed up on six days over 15 days total. Twenty-four hours before every imaging session, mice were injected with αCD35-iFlour647 i.v. to visualize the light zone (follicular dendritic cells (FDCs)) of GCs. Macrophages are visible due to their autofluorescence (pseudo colored in cyan and white). Example micrographs were acquired at λ_Ex_ = 840 nm and show autoreactive GC remodeling during initiation. Selected planes represent comparable focal planes. White arrows point to the same collagen trabecula to illustrate comparability. Note that tissue changes can only be visualized in 3D. Scale bar: 100 µm. (**E**) To investigate the contraction of autoreactive GCs, mice were treated with R848 for 4 weeks. Then the R848 treatment was stopped, and an AIW was implanted to observe the GC contraction on six days over 15 days total. Labeling and representative micrographs were selected and are annotated as in D. For representative purposes Top-hat and median filter (Despeckle) were applied in Fiji to enhance morphological features in all micrographs in this figure. Micrographs were furthermore enhanced in brightness and contrast (linear).

GCs were identified based on their FDC network (identified by intravital labeling for CD35/complement receptor 1 (CR1) (αCD35-iFlour647)) and the presence of autofluorescent vacuolar TBMs (*17, 49*). PT-tracks and the PALS were visualized by second harmonics generation (SHG) and dense GFP+ populations, respectively. Individual GCs underwent serial intravital microscopy and had to be reidentified every imaging day despite significant tissue changes (Fig. 2D and E). Therefore, a 3D overview, 250 µm deep (starting from the spleen capsule), of the spleen area within the window’s field of view, was acquired at the beginning of each imaging session to highlight specific patterns of multiple FDC networks (light zone of GCs) (Fig. 3A and B). In the first imaging session, the most superficial GCs were selected to be areas of interest. In the following imaging sessions, the selected GCs were reidentified in 3D overviews based on surrounding landmarks such as vasculature (lack of fluorescent emission surrounded by autofluorescent red pulp macrophages), other GCs, the capsule, or the borders of the window’s field of view (Fig. 3B). When setting the GCs of interest in focus, collagen trabeculae, visualized by SHG signal, were used to reposition the GC into the same imaging position as the previous imaging day by reidentifying and ‘aligning’ the collagen fibers in relation to the border of the field of view (FOV) (Fig. 2D+E).

**Fig. 3.**
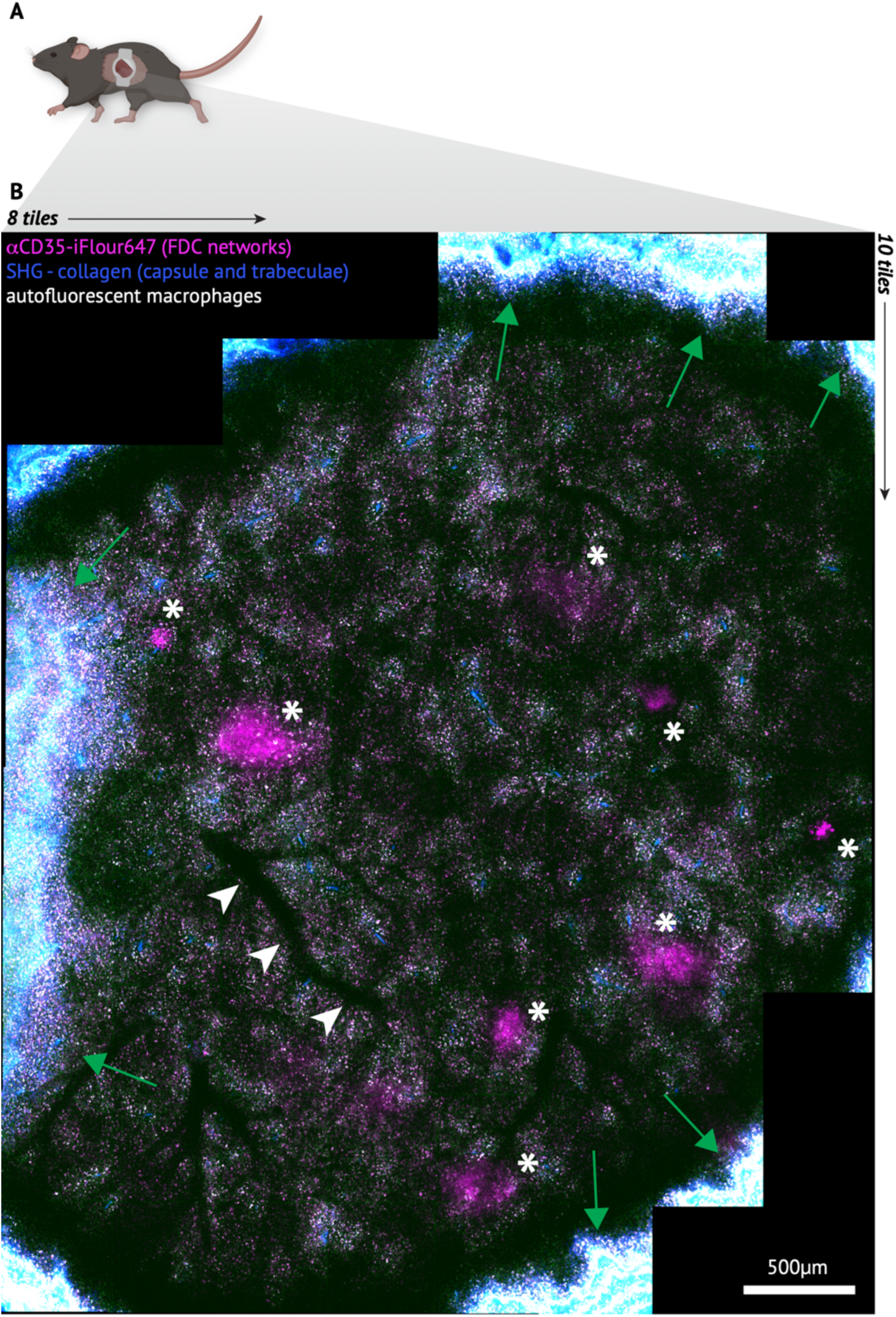
Overview of spleen within the entire AIW’s field of view. (**A**) Mouse with an AIW allowing serial intravital microscopy of the spleen. (**B**) 3D-stitched overview of the entire AIW’s field of view (8×10 tiles), created using Multi-area Time Lapse (MATL) function and Stitching Function in Fluoview while live imaging (acquired at λ_Ex_ = 840 nm). Here, the 3D overview is displayed as a Maximum Z-projection for representative purposes of the deepest 150 µm acquired (100-250 µm below capsule) and the micrograph is furthermore processed by subtracting background, median filtering (Despeckle) and adjusting linear brightness and contrast using Fiji. Intravital labeling with αCD35-iFlour647 visualizes the light zone (follicular dendritic cells (FDCs)) of GCs. Asterisks (*) point to most superficial FDC networks. White arrowheads point to a large blood vessel, used as landmark for reorientation over time. Green arrows indicate the borders of the window’s field of view. Scale bar: 500 µm.

### Follicles and their FDC network remodel extensively during GC expansion and contraction

To visualize 3D tissue remodeling in an easy-to-display 2D image, 3D imaging stacks were resliced orthogonally. The orthogonal view revealed that the labeled FDC network and the follicle expanded during R848 treatment (visualization in micrograph: FDC + “dark” areas scaffolded by autofluorescent macrophages) (Fig. 4A). Conversely, the FDC network and follicle were observed to contract after R848 treatment cessation (Fig. 4B). During GC initiation, non-labeled cells accumulated above the FDC network (“dark” area above FDC network) (Fig. 4A, Day 20). These “dark” areas were found to push the FDC networks of GCs deeper into the tissue as the R848 treatment progressed (Fig. 2D and Fig. 4A). This was most likely due to expanding proliferating B cells (non-labeled cells). In addition to proliferating dark zone B cells, also naive mantle zone B cells are unlabeled. Macrophage populations expanding in the red pulp above the follicles would be highly autofluorescent. Still, the accumulation of dark cells above the FDC network led to a higher degree of light scattering and thus decreased imaging quality (Fig. 2D and Fig. 4A). This complicated gaining true 3D imaging readouts, because areas of interest moved out of sight over time in the initiation cohort. We therefore excluded Day 20 from FDC analyses, as the FDC network seemed to disappear to non-imageable depths. Orthogonal views showed that the FDC network expanded during R848 treatment (Fig. 4C), whereas it contracted following R848 cessation (Fig. 4D).

**Fig. 4.**
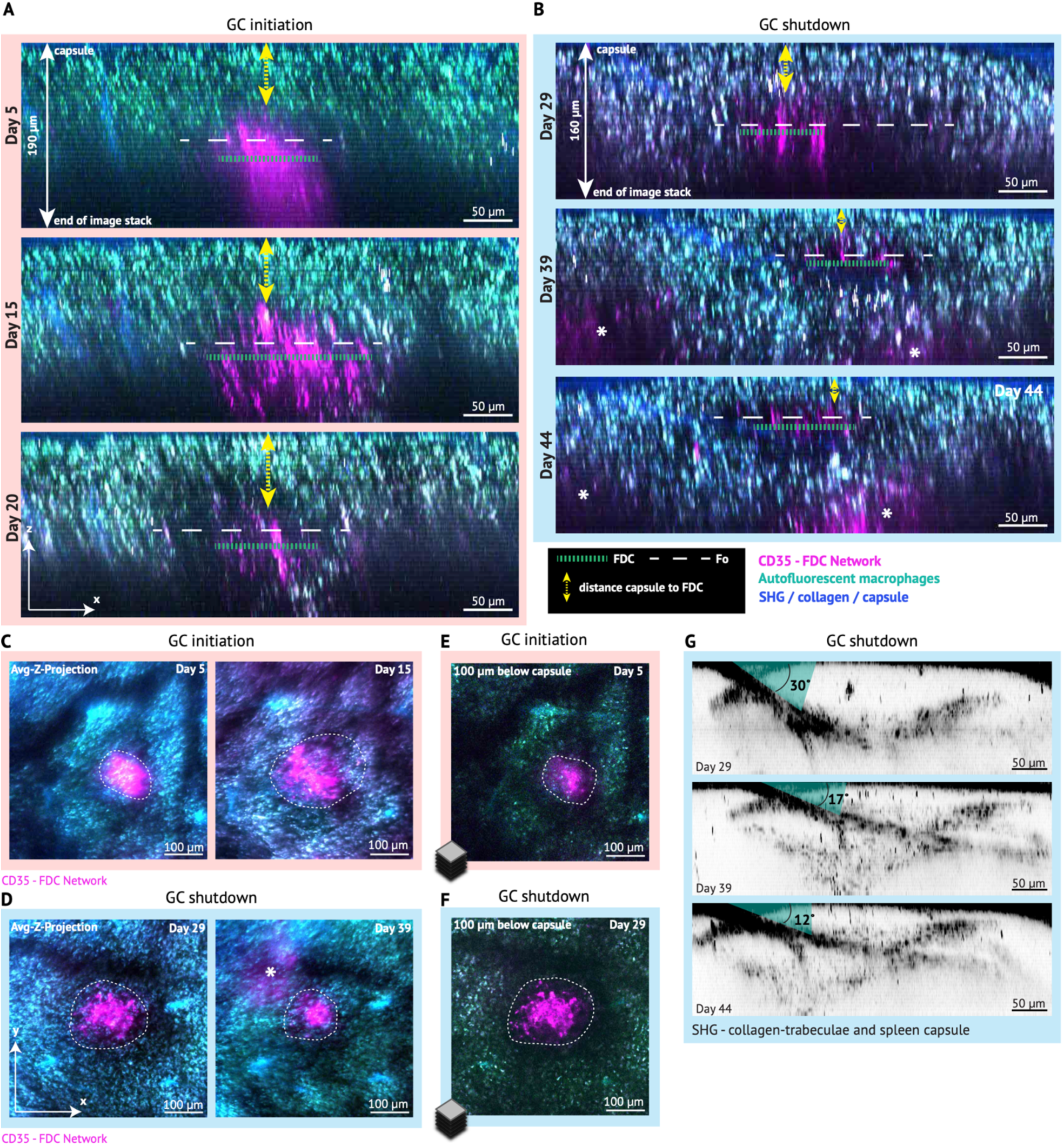
Image analysis of serially tracked splenic GCs. (**A**) Orthogonal view of autoreactive GCs during initiation over 2 weeks across 190 µm depth at all days (Day 5-20). The horizontal green dashed line indicates the diameter of the FDC network. The horizontal white dashed line indicates the estimated diameter of the follicle (Fo) (“dark” area). The yellow dashed vertical double-arrow line indicates the changes in distance from capsule to beginning of the FDC network. All images were acquired at λ_Ex_ = 840 nm, with a step size of 5 µm. Micrographs display Maximum Z-projection of all imaging frames with positive CD35 staining (intravitally labeled). Further information on Z-projections can be found in Fig. S1). Scale bar: 50 µm. (**B**) As A, but for the contraction phase (Day 29-44) and across 160 µm depth at all days. Note that deeper GCs are rising up from below over time (*) on day 39 and 44 as the general spleen parenchyma contracts. (**C**) Average Z-projection of all imaging frames with CD35 staining (intravitally labeled) of autoreactive GCs during initiation on Day 5 and Day 15 (further information on Z-projections can be found in Fig. S1). White dashed circle indicates the labeled area of the FDC network. All images were acquired at λ_Ex_ = 840 nm. Scale bar: 100 µm. (**D**) As C, but for the contraction phase. Note the rising up of deeper FDC networks over time (*) on day 39. (**E**) Selected representative frame of a 3D image stack of autoreactive GCs on the first observation day during GC initiation. White dashed circle indicates the labeled area of the FDC network. Scale bar: 100 µm. (**F**) As E, but for the contraction phase. (**G**) Visualization of spleen parenchyma remodeling and tissue contraction during GC shutdown, by tracking changes of the splenic collagen trabeculae over time while GCs are contracting. All images were acquired at λ_Ex_ = 940 nm. Micrographs are complementary to the micrographs in panel B) and pre-processed similarly. Furthermore, channel with second harmonic generation signal (SHG) was isolated and the LUT inverted to visualize SHG/collagen signal in black (capsule and trabeculae). Scale bar: 50 µm. All micrographs in this figure were adjusted in brightness and contrast (linear) for representative purposes.

Despite the slow FDC network disappearance in depth, the remodeling of the FDC network could be visualized and later on quantified using 2D average Z-projections of the FDC network as well (Fig. S1A and B, analysis shown later in Fig. 6). To determine whether choosing a Z-projection was a valid approach to measure the FDC network remodeling in only 2 dimensions per time point, we visually compared the Z-projection to a single Z-slice from the 3D imaging stacks. There was agreement between the FDC network outlines as shown by comparison of the first imaging day of both GC initiation (Day 5) and GC shutdown cohort (Day 29) (Fig. 4E and F, respectively).

Our observations further indicated that the entire spleen parenchyma, including the red pulp, adapted to the changes in follicle size. To investigate this further, we tracked changes in the arrangement of collagen trabeculae in the red pulp surrounding follicles (Fig. 4G). The angle of the trabeculae emerging from the capsule (Fig. 4G) was getting smaller over time as the GC was contracting, indicating that the collagen trabeculae are intimately associated with follicular structures.

Taken together, using the AIW with a combination of different intravital labeling techniques and analyses can reveal unique longitudinal information of immunological processes in the spleen including remodeling of GCs and the entire spleen parenchyma.

### Reporters capture single cells and structures of interest in deep spleen tissue over time

We used the *Foxp3*^DTR-GFP^ fluorescent transgenic reporter to visualize T_REG_ cells, especially those localizing to GCs, anatomically marking them T_FR_ cells (*50*) (Fig. 5A to I). Two-photon excitation enables deep tissue imaging (*51, 52*). However, image resolution is limited due to high scattering in deeper tissues (*52, 53*). GCs, the objects of interest, are mostly found in relatively high penetration depths (>100 µm below capsule). T_FR_ cells are localized equally deep and emit smaller and dimmer spots of signal than FDCs. To visualize T_FR_ cells reliably in all imaging depths, processing of images was needed to enhance contrast or signal to noise ratio (SNR).

**Fig. 5.**
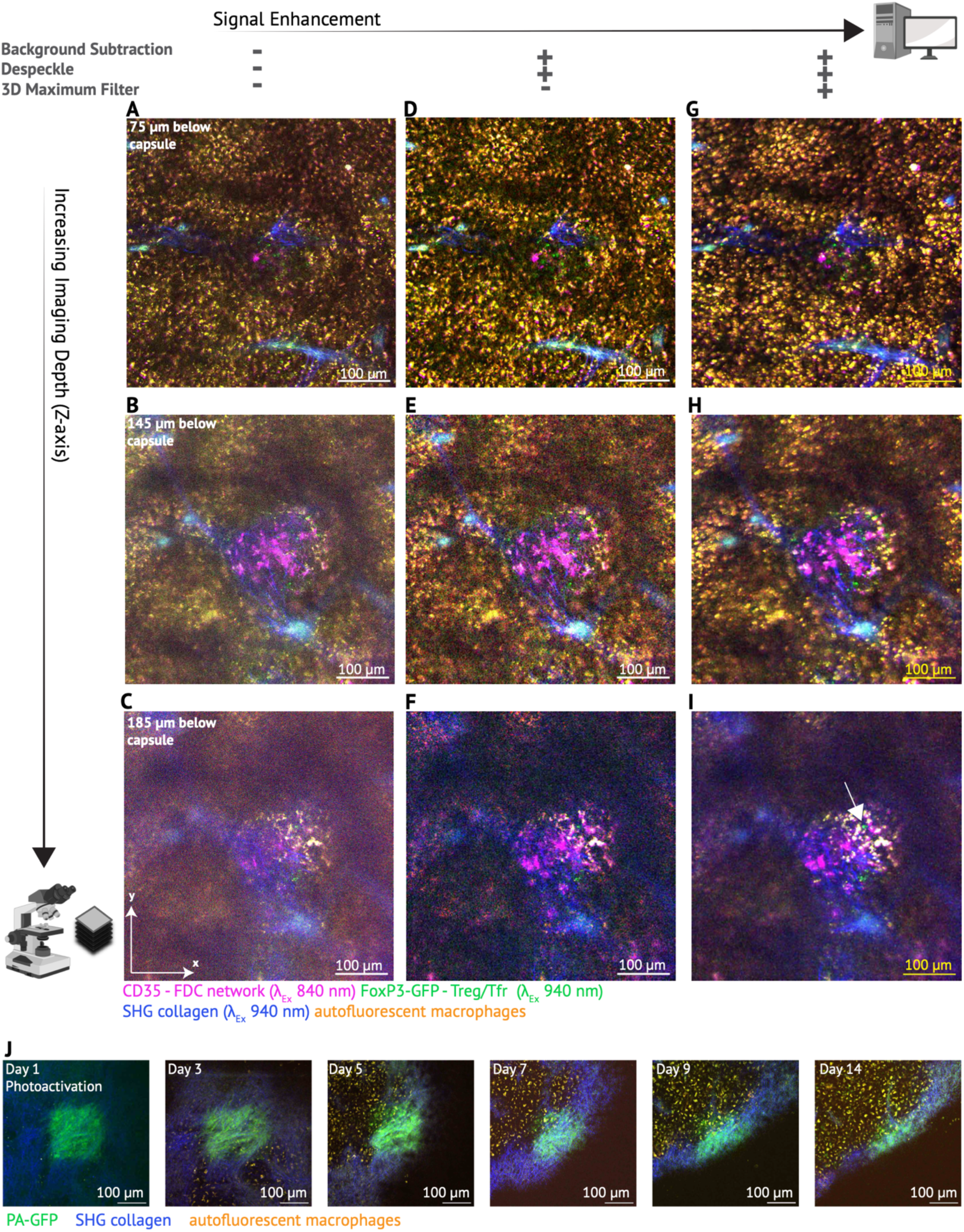
Image processing of deep and longitudinal 3D intravital spleen imaging data sets to enhance single cell signals. (**A-C**) Different focal planes of the same 3D imaging stack are displayed in overlay of both excitation tracks λ_Ex_ 840 and λ_Ex_ 940. Displayed imaging frames stem from 75 µm, 145 µm, and 185 µm below capsule of the same FOV. FoxP3-GFP+ cells are generally visible, but low signal-to-noise ratio (SNR) severely impedes manual cell counting. In A-C brightness and contrast were enhanced (linear), but no filter was applied. Scale bar: 100 µm. (**D-F**) Same images as A-C, additionally images were processed using Background subtraction and a median filter (Despeckle). FoxP3-GFP+ cell visibility improved but was occasionally still difficult to identify in deeper imaging planes. Scale bar: 100 µm. (**G-I**) Same images as D-F, with additional application of a 3D Maximum filter to enhance and spatially enlarge dim signal spots such as FoxP3-GFP+ cells throughout the entire 3D imaging stack. Note how GFP+ cells pop out bigger and brighter and are hence easier to quantify manually throughout different imaging planes of the entire 3D stack. The white arrow identifies one example FoxP3+ cell. Scale bar: 100 µm, calculated by pixels as precise scaling of image was not applicable after 3D maxima filter application due to spatial enlargement of single spots. (**J**) Longitudinal tracking of PA-GFP stimulated area for 2 weeks, acquired at λ_Ex_ = 940 nm. PA-GFP is visible the entire time, however field of view is slowly detaching during the last 10 days and rotating slowly out of sight. Slow detachment of fields of view is a typical technical challenge of serial intravital imaging of abdominal organs. Images are median filtered (Despeckle). Scale bar: 100 µm.

To reduce noise and background signal, we utilized easy-to-apply and open-source processing tools such as the “Despeckle” function and “Rolling Ball Background Subtraction”, both available in Fiji (*54*). We furthermore used different filters, enhancing maxima, to increase both size and signal of small, dim features such as T_FR_ cells in deep tissue. The effects of different (3D) filters on dim GFP emission from T_FR_ cells and the FDC network, are displayed in Fig. 5A-I at three different tissue depths.

We furthermore tested the longevity of photoactivatable (PA)-GFP in the splenic capsule (using B6.Cg-Gt(ROSA)^26Sortm1(CAG-PA-GFP)Rmpl^/J mice) and showed that PA-GFP can be tracked for 2 weeks after photoactivating an area of the splenic capsule (Fig. 5J). This allows longitudinal tracking of areas of interest in the spleen over long time periods.

### Microscopic GC dynamics correlate to macroscopic spleen dynamics

To begin to validate the method, we next examined whether the microscopically detected changes of the FDC network over time correlate to macroscopic changes in spleen weight in matched cohort studies. Splenomegaly, an enlarged spleen, is a hallmark of SLE (*55*) and is known to be triggered by R848 treatment (*48, 56*). Splenomegaly was detected 20 and 29 days into R848 treatment, then gradually reverted after the cessation of R848 treatment (Day 29 – 44) (Fig. 6A). The size was significantly different between untreated mice and all time points during R848 treatment and shortly after R848 cessation. Only on Day 44, the last observation timepoint after R848 treatment cessation, the size difference was not significant anymore (Fig. 6A).

**Fig. 6.**
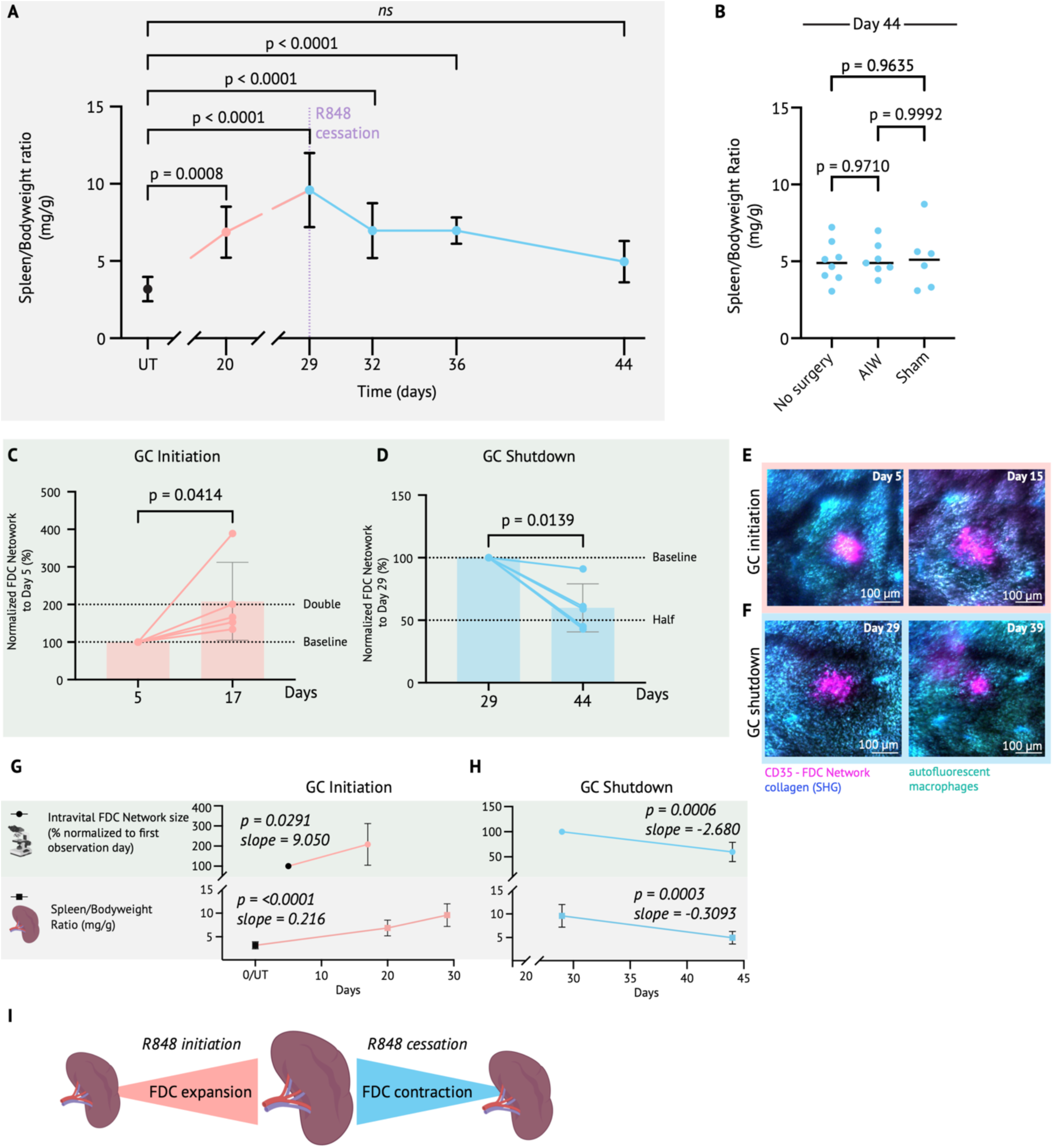
Macroscopic remodeling of the spleen correlates with microscopic remodeling of germinal centers. (**A**) Macroscopic spleen remodeling was evaluated using spleen to bodyweight ratio. Untreated (UT) mice, mice undergoing the R848 initiation regimen until day 20 or day 29, and mice undergoing the R848 cessation regimen until day 32, 36 or 44. Note that Day 20 was measured on AIW mice, all other time points were measured on mice that did not undergo any surgery. Experimental n = 4-11 from 2-4 cohorts per timepoint. The data are represented as means ± SD. P values were computed with ordinary one-way-ANOVA, comparing all time points to untreated control mice without an abdominal imaging window. (**B**) Comparison of spleen to bodyweight ratios between mice that did not undergo any kind of surgery, mice that were implanted with an AIW, and mice that underwent Sham surgery. All mice were in the R848 cessation group, organ harvest always on day 44. Experimental n = 6-8 mice per group. Data was pooled from 2-5 independent cohorts per group. (**C**) Measurement of the FDC area based on average Z-projections on the beginning (day 5) and day 17 of the GC initiation period (n = 6 from 3 mice of 2 cohorts; on day 17, 1 exclusion for technical reasons). (**D**) Measurement of the FDC area on average Z-projections during GC shutdown (n = 6 from 3 mice of 3 cohorts; Day 12, 1 exclusion for technical reasons). In C and D, the data are represented as individual measurements normalized to the baseline measurement on day 5 (initiation) and day 29 (cessation), respectively (initial measurements set as 100%). Means are indicated for each group. P values were computed on non-normalized, raw measured area values (Fig. S2) with two-tailed parametric paired t-test. (**E**) Representative images (average Z-projection) of the expansion of the FDC network during R848 treatment initiation, acquired at λ_Ex_ = 840 nm. Micrographs display Z-projections of all frames with positive CD35 labeling, median filtered and adjusted in brightness and contrast (linear). (**F**) As E, but for the contraction phase. **(G**) Comparison of trends in FDC network changes (green background) and macroscopic spleen remodeling (grey background) during GC initiation. P values were calculated using Pearson correlation between time (in days) and the microscopic and macroscopic metric readouts for each dataset. Slopes were calculated with simple linear regression analysis of each dataset. (**H**) As G, but for the contraction phase. (**I**) Graphical summary of correlation of microscopic remodeling detected by intravital imaging to macroscopic spleen remodeling.

Importantly, the observed spleen contraction after R848 treatment cessation on Day 44 was comparable between AIW implanted mice, Sham operated mice, and mice that did not undergo any surgical intervention (Fig. 6B). Furthermore, the spleen to bodyweight ratio on Day 20 was measured on mice that had an AIW implanted for 2 weeks during spleen expansion (Fig. 6A). Together, this confirms that both expansion and contraction of the spleen is possible in the presence of an implanted AIW over the spleen.

Microscopically, we found that the FDC network expanded to double its baseline size from Day 5 to Day 17 (Fig. 6C and E, and Fig. S2A). This trend correlated to the general spleen expansion (Fig. 6G). Conversely, the reversion of splenomegaly correlated with the FDC size reduction from Day 29 – Day 44 to almost half from baseline measurements on the first day of R848 cessation (Fig. 6D, F and H, and Fig. S2B).

In conclusion, the microscopic expansion and contraction of the FDC network could be correlated to, and thereby validated by, macroscopic spleen remodeling. This demonstrates that macroscopic dynamic changes are mirrored in long-term intravital dynamic microscopic readouts of the spleen through an AIW, such as remodeling of the FDC network (Fig. 6G to I).

### Serial intravital 2-photon imaging of the spleen aligns with flow cytometry read-outs

We furthermore correlated the intravitally detected microscopic FDC network changes during the GC contraction phase (Fig. 6D and F) with flow cytometry readouts from matched cohort mice on respective imaging days (Fig. 7). R848 treated mice were matched both on Day 29 (beginning of observation of GC contraction) and Day 44 (end of observation of GC contraction) to determine if intravitally observed changes associated with GC contraction are reproduced in flow cytometry read-outs. Additionally, we included matched untreated controls. We furthermore compared Day 44 in non-operated mice, sham-operated mice and AIW-operated mice to further investigate the influence of surgery, window implant, and imaging on flow cytometry read-outs. Finally, we divided the spleen of AIW mice into the part that was attached to the AIW and the part that was distal from the AIW, to investigate if the window attachment influenced the underlying tissue (Fig. 7A). The gating strategies can be found in Fig. S3. The flow cytometry analyses showed that after a transient peak on Day 29, GC B-cell frequencies across non-surgery and surgery mice (both Sham and AIW) had fallen to approximately the same levels on Day 44 yet remained above the level of untreated mice (Fig. 7B). There were no significant differences between unoperated and operated mice with or without an AIW at Day 44 (Fig. 7B). T_FH_ cell frequencies also increased during R848 treatment and slightly decreased after R848 treatment cessation. The decrease in T_FH_ cells in AIW mice (both in window-proximal and distal spleen parts) seemed to be slightly stronger than in Sham and unoperated mice (Fig. 7C).

**Fig. 7.**
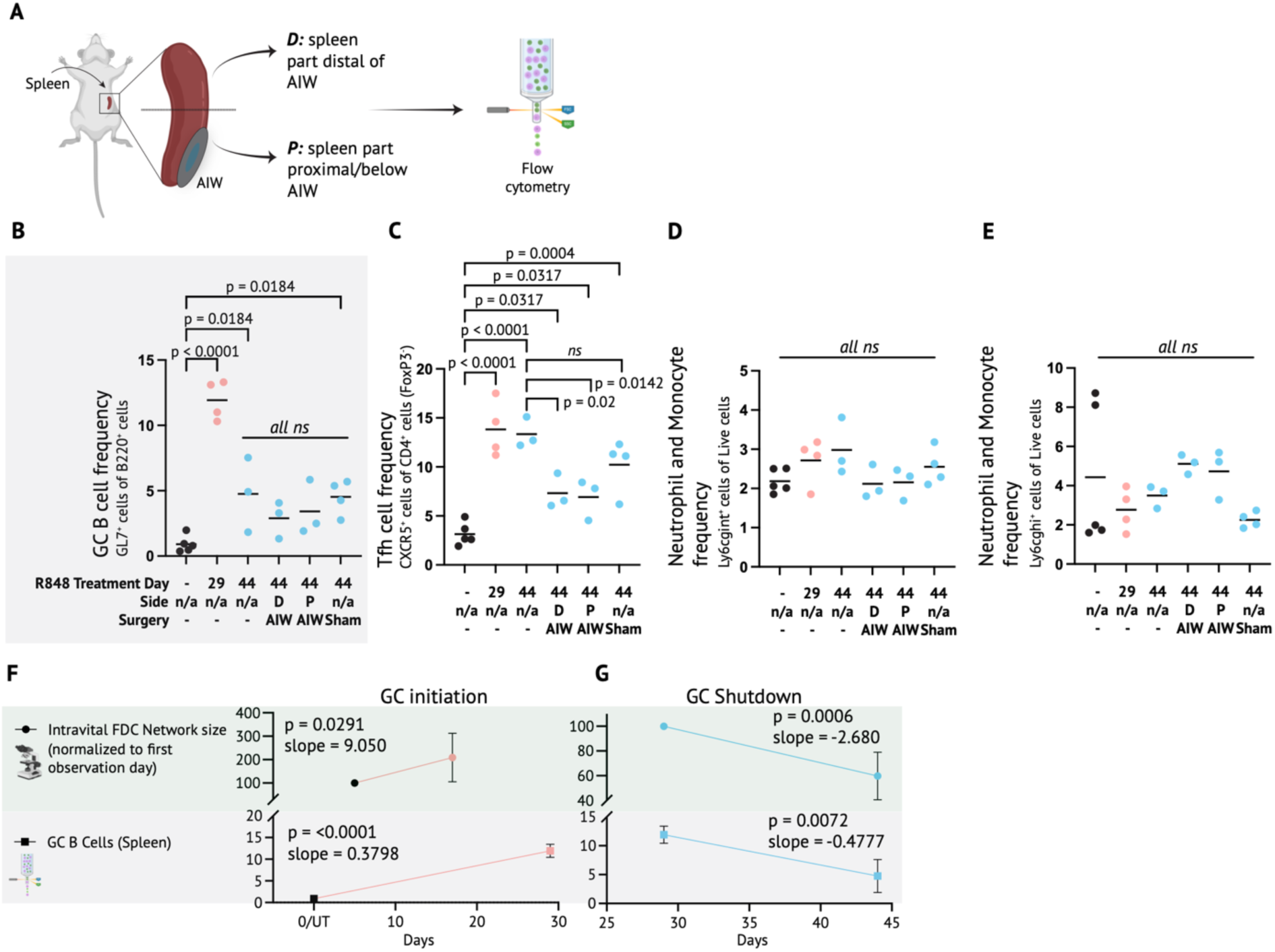
Serial intravital imaging of the spleen agrees with and complements flow cytometry analyses. (**A**) Schematic overview of sampling for flow cytometry: Spleens from AIW mice were divided into a part that was attached to the AIW (proximal) and a part that was not in direct contact with the AIW (distal). Spleens from Sham-operated or non-operated mice were not divided in this manner, but an equally big spleen piece was processed. **(B-E)** Experimental n = 3-5 from 1-2 cohorts at different days of the R848 treatment regimen. Individual datapoints are represented with the mean. P values were computed with one-way ANOVA (GCB, T_FH_) or Kruskal-Wallis’s test (Ly6gc) comparing all groups to untreated controls. Furthermore, surgery mice and non-surgery mice were compared at Day 44. **(B**) Flow cytometry analysis of GCB frequencies in the spleen. **(C**) Flow cytometry analysis of T_FH_ frequencies in the spleen. (**D**) Flow cytometry analysis of Ly6gc^int^ frequencies in the spleen. (**E**) Flow cytometry analysis of Ly6gc^hi^ frequencies in the spleen. (**F**) Comparison of the trends in the FDC network changes and GC B-cell frequencies during GC initiation. (**G**) As F, but for the GC contraction phase. P-values were calculated with Pearson Correlation of time (days) vs. each dataset, respectively. Slopes were calculated with simple linear regression analysis of each dataset.

Furthermore, we found that there was no major inflammatory response associated with the AIW implant itself, neither in the part of the spleen that was attached to the window, nor the part that was distal to the AIW (Fig. 7D and E). Whereas Ly6cg^int^ frequencies where similar, AIW mice seemed to have slightly higher Ly6cg^hi^ frequencies than unoperated or sham operated mice at day 44, but when comparing to the untreated controls, there was no significant difference (Fig. 7D and E).

Interestingly, GC B-cell levels could be correlated to changes in the FDC network observed microscopically and intravitally over time, both increasing during R848 treatment and decreasing following R848 cessation (Fig. 7F and G). Summarizing these observations, the observed FDC expansions (Day 5 until Day 20) and contractions (Day 29 until Day 44) intravitally, were confirmed by flow cytometric read-outs, especially by the similar trend of GC B cell levels over a comparable time frame (Fig. 7F and G). Overall, we confirmed that our intravital findings correlated well with flow cytometric readouts, as both FDCs and GC B cells are pivotal GC cell populations. We furthermore excluded any major inflammatory influence based on surgical intervention or chronic AIW implantation. This both validated the immunophenotype readout acquired through serial intravital microscopy upon the different R848 treatment regimens and served as a further validation that the AIW did not perturb the immune response.

### 2-photon imaging of spleen explants complements serial intravital imaging

We previously noted a “dark” area surrounding the labeled FDC network in serial intravital images (dark area lacking abundant autofluorescent macrophages). This dark area expanded during R848 treatment and contracted upon R848 treatment cessation (Fig. 4A and B). We assumed that the dark area marked both the mantle zone and the biological dark zones of GCs (proliferating and affinity-maturing B cells). To test this hypothesis, we outlined the dark area next to the FDC network in freshly explanted spleens and later stained the same areas for the proliferation marker Ki67 (Fig. 8A to E). Additional to the usual intravital labeling of αCD35-iFluor647, we added the intravital label CD169-PE before euthanasia to mark metallophilic macrophages of the marginal zone (Fig. 8F). After imaging the freshly prepared spleen sections, they were fixed and stained with αKi67-A488 (Fig. 8G). This sequential imaging approach was necessary to visualize the “dark” areas as closely as possible to the intravital images and because the subsequent fixation and permeabilization process eliminated the GFP (FoxP3-DTR-GFP) signal and decreased the signal for the intravital labeling of αCD35 and αCD169. To achieve optimal overlay results for the sequentially imaged area, a precise documentation and reproduction of the spleen slice position on coverslips and a final digital image registration was performed (Fig. 8A-E). This allowed us to associate the intravitally detected “dark” areas (Fig. 8F+H) to the actual biological dark zone of GCs (Fig. 8G+I). It furthermore validated the approach of localizing GCs by outlining FDC networks and TBMs (Fig. 8H), the phagocytes of active GCs (*57*). TBMs are bigger and more vacuolar than regular macrophages and are thus readily distinguishable from other macrophages.

**Fig. 8.**
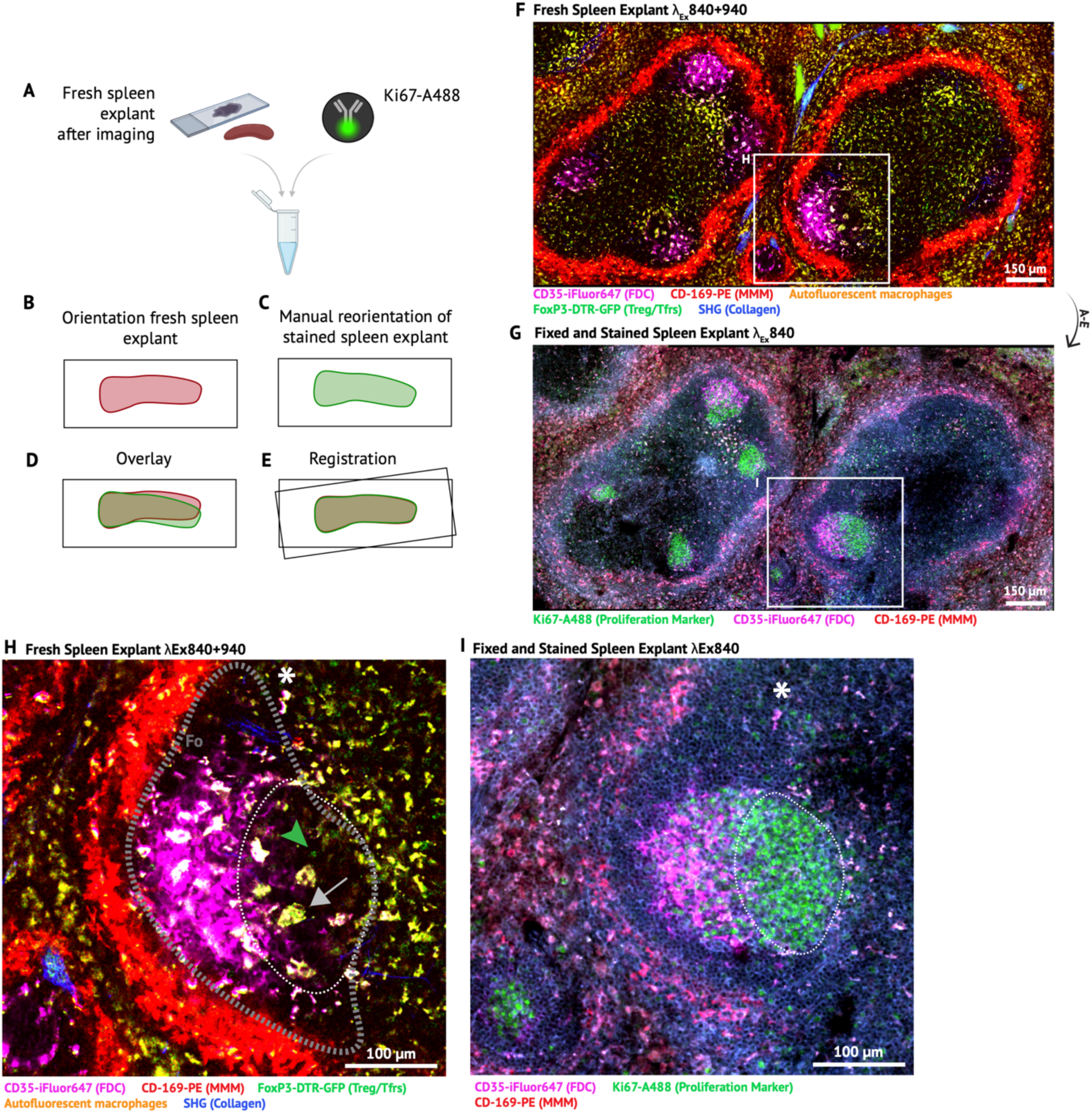
Spleen explant imaging confirms and complements intravital imaging readouts. (**A-E**) Schematic overview of tissue handling and image processing to ensure identification of the same areas imaged in a freshly prepared spleen explant and after tissue fixation and staining with the proliferation marker Ki67. The spleen slices are stained free-floating in a 1.5 ml tube. Therefore, orientation can easily be lost if not documented properly (A+B). (**F**) Overview of two white pulp areas of a fresh spleen explant, showing several FDC networks (CD35). Labeling strategies and pseudo-coloring indicated in legend. Scale Bar: 150 µm. (**G**) Same area overview as F, after tissue fixation and staining with Ki67-A488 to visualize proliferating cells. Labeling strategies and pseudo-coloring indicated in legend. Scale Bar: 150 µm. (**H+I**) Cut-out and magnification of the same areas of F+G, respectively. Literally “dark” areas of freshly explanted spleens are outlined (dotted circle in white and additional CD35-negative area of the follicle (Fo)). These “dark” areas correlate partly to the biological dark zone of GCs (marked by proliferation marker, indicating proliferating GC B cells). The remaining “dark” area of the respective follicles is assumed to be the mantle zone. The white asterisk (*) points to the same macrophage, indicating the same focal plane is in focus in the fresh (H) and fixed (I) spleen section. Labeling strategies and pseudo-coloring are indicated in legend. Scale Bar: 100 µm. In H, the grey arrow points to a GC macrophage (tingible body macrophage, TBM), and the green arrowhead points to a FoxP3-DTR-GFP^+^ cell in the GC, here a T_FR_ cell. All micrographs in this figure were acquired in MATL-Tile-Scan mode and are stitched (4×2 Tiles). For representative purposes, stitched micrographs were median filtered (Despeckle) and adjusted in brightness and contrast (linear). Excitation wavelengths indicated in figure.

### Mice thrive with an implanted AIW

Apart from verifying the biological readouts gained by serial intravital imaging, we furthermore aimed to examine the impact of the AIW on the well-being of the mice. With correct placement of implant between spine, ribcage and hind leg, we found that mice with an AIW over the spleen, were able to move and climb freely in their cages without any restrictions in movement (Fig. 9A). They did not show any signs of distress or discomfort based on daily monitoring. To furthermore objectify these observations, their bodyweight was measured every day at the same time during the entire experiment. To ensure precision, AIWs were weighed before implantation and the weight was subtracted from the measured mouse body weight after implantation. The results demonstrated that all mice with an AIW reacted solely with a transient mild weight loss to the surgery and first imaging sessions (Fig. 9B and C). Mice undergoing R848 treatment initiation showed a significant weight loss, albeit less than 10% of body weight, after surgery (Fig. 9B). Mice undergoing R848 treatment cessation in conjunction with AIW implantation only lost approximately 5% of body weight, even though at the time of surgery mice still received one last R848 treatment (Fig. 9C). Neither group lost any further weight after the weight had normalized and stabilized approximately 1 week post-surgery (Fig. 9B and C). Mice undergoing sham surgery did not show any significant changes in body weight (Fig. 9D).

**Fig. 9.**
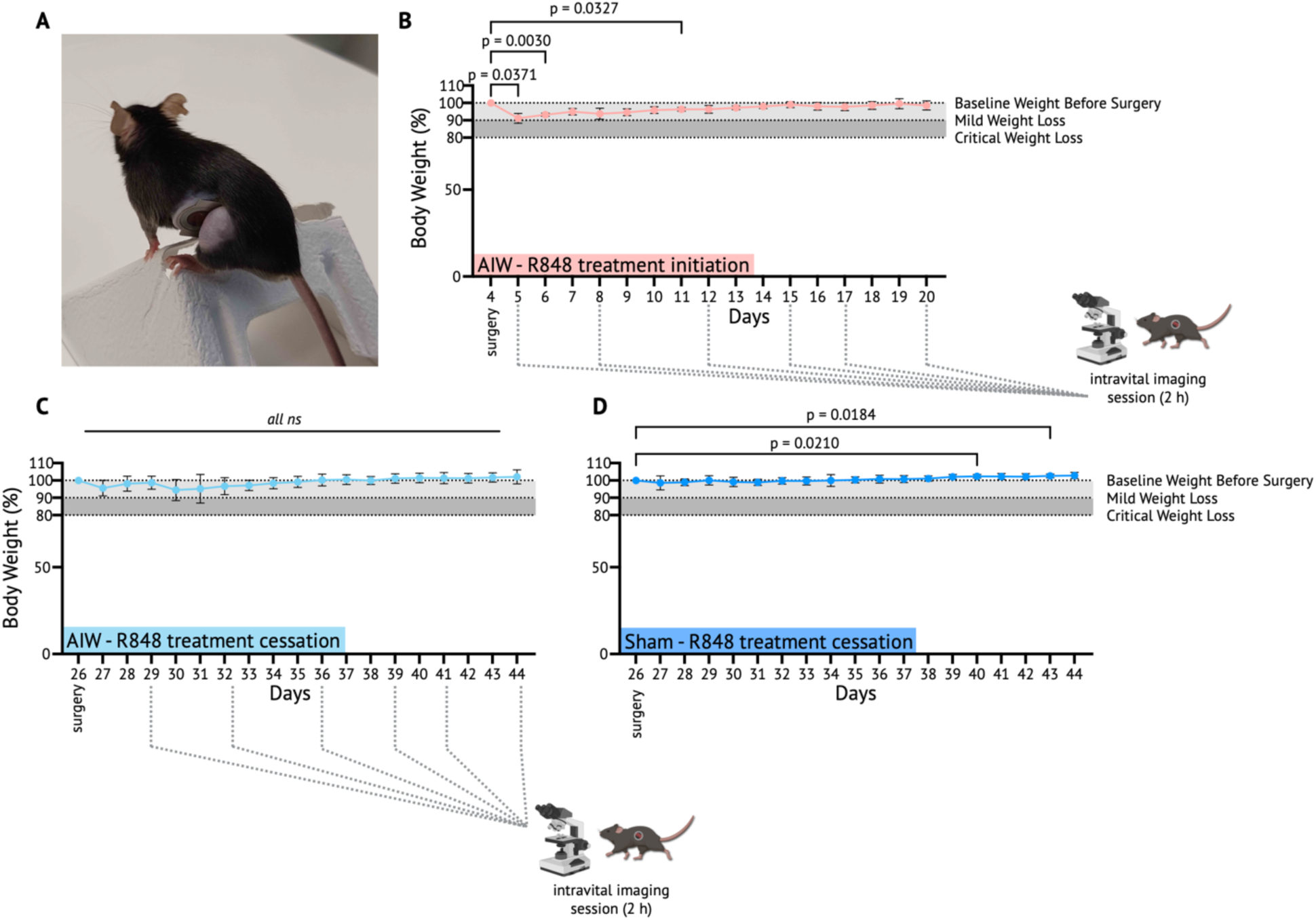
An AIW does not impact the thriving of mice. (**A**) Mouse, 6 days after AIW implantation. The mouse is able to climb as usual when the AIW is correctly placed between spine, ribcage and hind leg. (**B**) Weight curves of mice with implanted AIW undergoing the R848 GC initiation treatment regimen (n = 4 from 2 cohorts). (**C**) Weight curves of mice with implanted AIW undergoing the R848 GC shutdown treatment regimen (n = 8 from 5 cohorts). (**D**) Weight curves of sham operated mice without an implanted AIW undergoing the R848 cessation regimen (n = 6 from 3 cohorts). Bodyweight curves of mice were normalized to the bodyweight of each individual mouse on surgery day. The weight of the implanted AIW was subtracted from the measured bodyweight after AIW implantation. Light and dark grey areas indicate weight loss of 10% (mild) or 20% (critical), respectively. Below the grey areas, a weight loss would be indicated as critical, >20%, marking a humane endpoint. The baseline (100%) for bodyweight was set on surgery day before the surgical procedure for each mouse individually. Grey dotted lines indicate intravital imaging days. Data are represented as means ± SD. Adjusted P-values were computed with ordinary one-way ANOVA for multiple comparisons. Only P < 0.05 are indicated in the graph.

### AIW implant over the spleen does not induce a systemic immune response

To further examine the biological impact of the AIW, we compared the skin-draining inguinal lymph nodes (IngLNs) on the non-surgery/contralateral side (right) and the surgery/ipsilateral side (left). We further investigated the influence of surgery in general and of the AIW specifically by comparing the IngLNs of sham, implant, and non-surgery control mice. We compared the weight of IngLNs for basic assessment, results of flow cytometry for a detailed immune cell phenotyping, and confocal microscopy to investigate anatomical changes in lymphatic vascularization (Fig. 10A). IngLNs were harvested at the end of experiment, 2 weeks after surgery (AIW and Sham operated mice) and after a total of 6 imaging sessions per mouse (AIW) (Fig. 10B to D). IngLNs from the R848 initiation group were harvested on Day 20, whereas IngLNs from the R848 cessation group were harvested on Day 44. Mice without surgical intervention from matched cohorts were euthanized at Day 44, the end of the R848 treatment cessation experiment, or were entirely untreated control mice (Fig. 10E and F). Already the initial IngLN weight analysis showed significant differences between the two sides in mice that underwent any kind of surgery (AIW and Sham surgery mice), the IngLN on the side of surgery (left) being significantly heavier in all 3 surgery groups (Fig. 10B to D). However, left IngLN sizes were comparable between AIW and Sham surgery groups. In mice not undergoing surgery, there was no difference between sides (Fig. 10E and F).

**Fig. 10.**
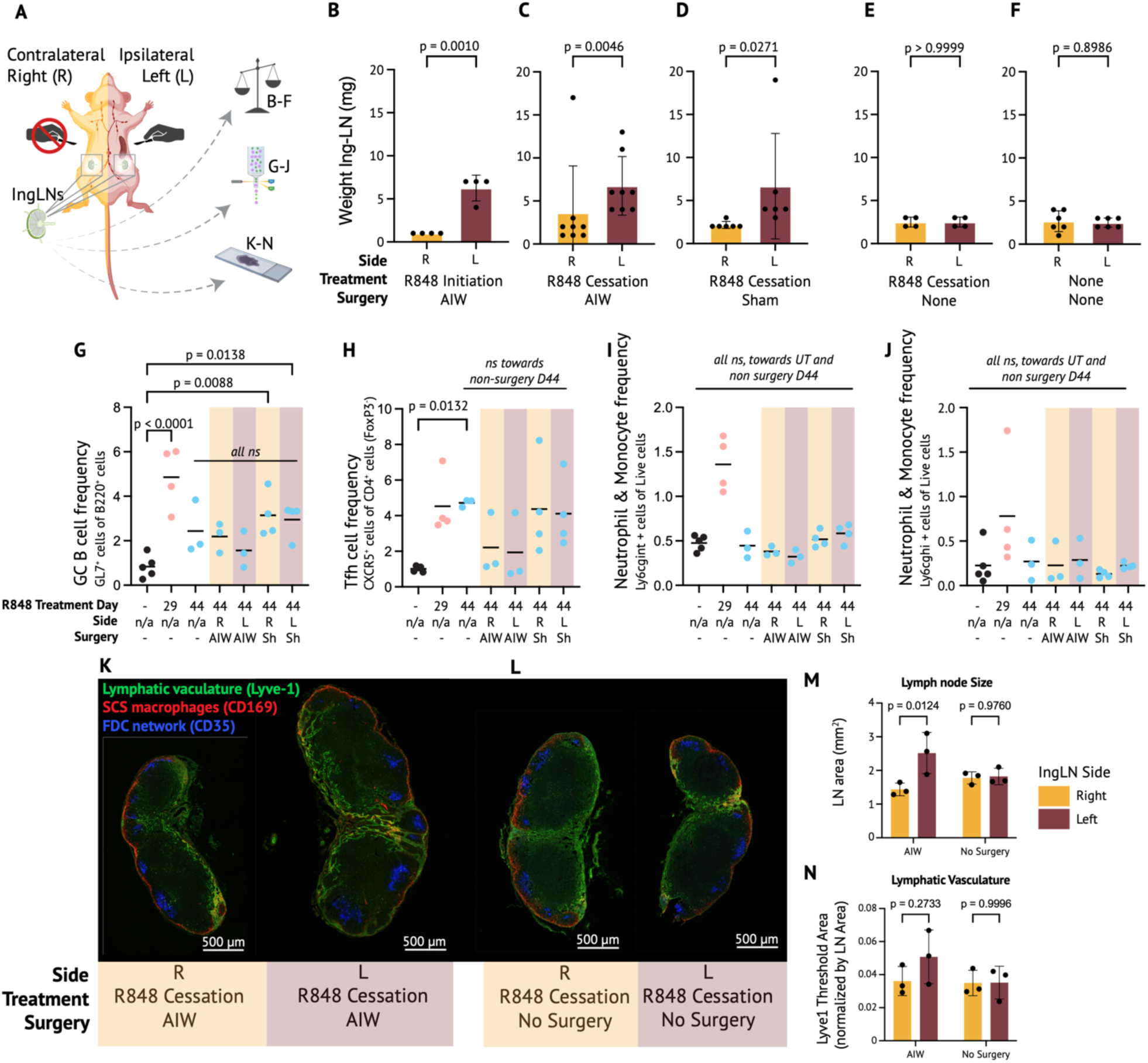
The AIW does not initiate a systemic immune response. (**A**) Schematic overview of experiment: Skin draining inguinal lymph nodes (IngLNs) from both ipsilateral surgery side (left - L) and contralateral non-surgery side (right - R) were harvested on last day of imaging experiment (2 – 2.5 weeks after surgery and 6 imaging sessions per mouse, Day 20 and Day 44). IngLNs were compared in weight, flow typing and microanatomical differences regarding their lymphatic vascularization. (**B**) IngLN weight of AIW mice that underwent GC initiation R848 treatment regimen (n = 4 from 2 cohorts). (**C**) IngLN weight of AIW mice that underwent GC shutdown R848 treatment regimen (n = 8 from 5 cohorts). (**D**) IngLN weight of Sham operated mice that underwent GC shutdown R848 treatment regimen (n = 6 from 3 cohorts). (**E**) IngLN weight of mice that underwent GC shutdown R848 treatment regimen, without any surgical intervention (n = 4 from 3 cohorts). (**F**) IngLN weight of untreated mice that underwent no surgical intervention (n = 6 from 2 cohorts). All weight comparisons of IngLNs are visualized in linear scale (unit: mg), data are represented as means ± SD, and statistical analysis was performed on lognormal transformed data. P-values were computed with two-tailed paired t-test. (**G**) Flow cytometry analyses of GC B-cell frequencies in IngLNs. (**H**) Flow cytometry analyses of T_FH_ frequencies in IngLNs. (**I**) Flow cytometry analyses of Ly6gc^int^ frequencies in IngLNs. (**J**) Flow cytometry analyses of Ly6gc^hi^ frequencies in IngLNs. Flow cytometry results of IngLN analyses stem from mice at different R848 treatment regimen time points, data from 3-5 mice per group (1-2 cohorts). Either both IngLNs were pooled, or right and left side separated. Individual datapoints are represented with mean. P-values were computed with Kruskal-Wallis’s test (GCB, T_FH_, Ly6gc) and compared to untreated (UT) controls. Furthermore, mice undergoing surgery were compared to non-surgery mice on day 44 as well. Only significant P-values shown. (**K**) Representative immunofluorescence confocal micrographs of right and left IngLN from a mouse with an implanted AIW and a mouse at day 44, stained with CD35-iFluor647 (blue), CD169-PE (red), and Lyve-1-eFluor450 (green). Scale bar: 500 µm. Micrographs are stitched confocal tile scans and adjusted in brightness and contrast (linear over entire stitched micrograph) (**L**) As K, but for a mouse without surgical intervention. (**M**) Quantitative image analysis of the IngLN size, measured by the IngLN area of each histological section (excluding fat or extranodal lymphatic vasculature). **(N)** Quantitative image analysis of the lymphatic vasculature (intranodal thresholded Lyve1+ area, normalized by general LN area. (M+N) n = 3 mice per group, all IngLN harvested from the R848 treatment cessation regimen groups on day 44. Areas of both IngLN section and thresholded Lyve1+ staining were measured using Fiji. Data are represented as means ± SD. P-values were determined using 2-way ANOVA, and post-hoc Šídák’s multiple comparisons test comparing right and left IngLN in each group. All P-values shown.

Flow analyses showed that the R848 treatment induces a robust GC response in IngLN (Fig. 10G and H, untreated vs. Day 29). This response was reverting in the R848 cessation cohort, though not reaching baseline levels again at Day 44. There was no significant difference in the GC response between unoperated mice, Sham or AIW operated mice on Day 44 (Fig. 10G and H). Moreover, there were no significant differences in neutrophil frequencies between control and Day 44 surgery groups (Fig. 10I and J). We observed, however, a transient influx of neutrophils at the peak of autoimmune inflammation at Day 29, which reverted to control levels on Day 44, 2 weeks after R848 cessation (Fig. 10I and J). When comparing contralateral (right) and ipsilateral (left) IngLNs of operated mice, no side differences were detected in any immune cell readout, except for a marginally higher Ly6cg^hi^ frequency in ipsilateral nodes of Sham-operated mice (Fig. 10G to J and Table S1). Flow gating strategies can be found in Fig. S3.

To follow up on the detected weight differences in contralateral and ipsilateral IngLNs of surgery mice, we performed immunofluorescence confocal microscopy of the lymphatic vasculature in IngLNs of AIW operated mice and unoperated mice at Day 44, both from the R848 cessation cohort. This revealed a notable increase in Lyve-1 expression of the ipsilateral IngLN of AIW-implanted mice (Fig. 10K) in comparison to IngLNs of unoperated mice (Fig. 10L). Generally, the ipsilateral IngLN of AIW-implanted mice was bigger (measured in area, Fig. 10K+M). Furthermore, the Lyve-1 positive area was increased in the ipsilateral IngLN of AIW-implanted mice (Fig. 10K), but this was not significant when normalized for IngLN area (Fig. 10N). Lyve-1 is a marker for lymphatic vasculature and the side-specific expansion of lymphatic vessels in skin-draining lymph nodes upon surgery may therefore contribute to the observed changes in weight (Fig. 10C) and size (Fig. 10M). These effects were only detectable in the ipsilateral IngLN (Fig. 10K, M, N) and did not seem to affect the contralateral IngLN (Fig. 10L-N)

Overall, these results indicate that the impact of both the surgery itself and the AIW implant is confined to the ipsilateral side, detected in weight (Fig. 10B to E), size (Fig. 10M), and discrete changes in lymphatic vasculature (Fig. 10K and L). Immune cell phenotyping showed no differences between operated mice and non-operated mice (Fig. 10G to J), as well as negligible side differences (Table S1). In summary, the AIW implant and serial intravital imaging of the spleen did not perturb the immune system systemically.

## Discussion

Previous studies using intravital microscopy of the spleen have uncovered several dynamic processes, such as the entry of T cells into T zones on perivascular pathways (*58*), the shuttling of B cells between compartments (*3*), or B cell retention in the marginal zone triggered by red blood cells (*59*). However, the terminal nature of the experimental setups employed in those studies prevented the capture of slower dynamics of longer-lasting splenic tissue changes. Ritsma and colleagues developed an AIW that allowed tracking the migration of intravenously transferred fluorescently labeled B and CD8+ T cells into the spleen. They then followed the acute CD8+ T cell response to antigen (ovalbumin peptide) challenge *in vivo* over multiple days and were able to quantify a T-cell response 7 days after antigen challenge (*11*). Here we show that serial intravital 2-photon microscopy using an AIW allows insight into week-long biological processes, exemplified by autoreactive GC dynamics (Fig. 2-4) and associated tissue changes (Fig. 2, 4, 6, and 7). This may allow novel high-dimensional temporospatial insights into important biological processes, such as the break-of-tolerance occurring in autoimmunity, vascular changes in disease, and hematological malignancies affecting the spleen.

Titanium, constituting the frame of the AIW, is recognized to be biocompatible (*60*). The glass coverslip can however be affected by cellular overgrowth, which can lead to reduced imaging quality, angiogenesis, and potentially an untoward immune response. This necessitated the implementation of an anti-fouling method to dampen the immunostimulatory activity of the window (*47*). A high-temperature PMOXA passivation was found to alleviate untoward immune reactions (Fig. 1), additionally prolonging the imaging period after surgical intervention by preventing organ encapsulation. Indeed, we found that the inflammatory stimulus of the initiation and cessation of R848 treatment is much stronger than any impact of the AIW itself, as measured by e.g. the GC B-cell response. This aligned well with our phenotyping of the immune environment in the spleen, where we found that the serial intravital imaging of GCs, including the surgical procedure, did not alter general immune responses nor trigger a systemic immune response (Figs. 7 and 10). Specifically, we compared contra- and ipsilateral skin-draining lymph nodes of the surgical area (Fig. 10) and spleen parts attached to the coverslip versus more distal to the AIW (Fig. 7). Local confinement of the inflammation to the ipsilateral (left) skin-draining IngLN was further proof that the AIW is not majorly perturbing the immune system and that any readouts through the window are not skewed by the implant.

After surgery, mice exhibited a slight and transient drop in bodyweight, (Fig. 9B and C). Repetitive 2-hour long anesthesia during each intravital imaging did not seem to affect the body weight long-term, potentially only the first imaging session induced a slight weight drop as mice had to adapt to the procedure initially. The ability of mice to thrive on this protocol is not only desirable from an ethical perspective but is also envisioned to deliver more reproducible results (*61, 62*). It is known that transient leukocyte trafficking into a surgical site decreases 72 hours post-surgery and is significantly increased by stress (*63*). Accordingly, in the R848 cessation cohort, mice were given 72 hours of rest after the surgical intervention before the seral intravital 2-photon observational period started. However, this was not possible for the R848 initiation cohort, which were only given 1 rest day due to technical challenges associated with the expansion of the spleen over time. Due to the affixture of the spleen to the window, the parenchyma cannot expand laterally, but only depth-wise, causing a more rapid decline in the imaging quality in depth (Fig. 2D and 4A), whereas in the cessation cohort the initial attachment of the spleen to the AIW was done on the already expanded spleen, preempting this issue. To be able to capitalize on imaging readouts as long as possible, it was necessary to implement a shorter rest period after surgery for GC initiation mice.

As the red pulp in the superficial areas of the spleen is extremely uniform without any specific landmarks, it was challenging to identify the same GC again and again over several weeks. A number of techniques have been described in the field of serial intravital imaging to reidentify previously imaged areas again, such as numbered gridded coverslips (*41, 64*), microcartography based on deposited microspheres (*65*), or laser-induced injury marks (*39*). We particularly refrained from the latter two as implanted microspheres and laser-induced injury marks might have perturbed the immune system. We found creating a 3D-stitched overview map in the beginning of every imaging session, capturing deep labeled FDC networks in the window area, to be a suitable strategy for reidentifying GCs in the spleen (Fig. 3). The overview map served to pinpoint the position of each GC in relation to the window position. The stable location of GCs relative to the architecture of especially rare large blood vessels further facilitated their reidentification (Fig. 2). This approach was reliable in identifying the same GCs over time, except when a FOV rotated out of sight, or the GC of interest was pushed too deep into the parenchyma due to heavy spleen remodeling as described above.

Through FDC labeling, we were able to identify the light zone of GCs, and further the entire GC based on the lack of red pulp macrophages in the dark zone next to the FDC area (Fig. 8F to I). We were hence able to capture both autoreactive GC expansion (Fig. 2D and Fig. 4A and C) and autoreactive GC contraction (Fig. 2E and Fig. 4B, D and G). We were also able to detect single cells of interest in the GCs (Fig. 5A to I). Despite being attached to the AIW, the spleen can remodel heavily, stays intact, and does not rupture while remodeling (Fig. 6A and B). In fact, the spleen could both expand and shrink with an AIW attached, as the spleen/bodyweight ratio doubled from baseline (untreated) to Day 20 (end of R848 initiation period) whereas it contracted to half its maximal size in just 2 weeks after R848 cessation (Day 29 to Day 44) (Fig. 6A and B). However, the extensive tissue remodeling, especially during the expansion of GCs caused a slow decrease of imaging quality over time as more tissue, potentially proliferating B cells, accumulated above the area of interest (Fig. 4A).

By comparing spleen metrics and flow cytometry results to serial intravital images, we confirmed that GC dynamics observed longitudinally intravitally (Fig. 2 and 4) correlated to developing splenomegaly and its reversion (Fig. 6) and to changes in flow typing results, such as GC B-cell frequencies of matched cohort studies (Fig. 7). These comparisons underline that the initiation and contraction of an autoreactive immune response can be captured intravitally. We furthermore show that intravitally detected changes in the light zone and the “darker” areas nearby can be indirectly linked to changes of the dark zone of GCs, as we associated the “dark” areas to the dark zone of GCs (Ki67) of spleen explants (Fig. 8). This demonstrates that longitudinal changes of the light zone mirror changes of the dark zone, making intravitally detected changes in FDC networks a correlate of changes in proliferating GC B-cell frequencies (Figs. 7 and 8).

While we will refrain from outlining technical challenges that most intravital imaging experiments have in common (*66, 67*) (exemplified in Fig. 5J), there are potential biological limitations that are specific to this method. Even though we significantly extended the imaging period to catch dynamic intravital remodeling, 2 weeks is still a narrow window in perspective of the general development of e.g., autoimmune diseases, in which autoantibodies often gradually increase for years in asymptomatic patients leading up to clinical diagnosis (*68*). Moreover, despite many similarities between the immune response in mouse and man (*69*), there are also many significant differences (*70*). Perhaps more importantly, the microanatomy of the spleen differs, mainly in the spatial arrangement of B- and T-cell zones (*71, 72*). That said, the functional spleen microanatomy is still poorly defined (*71*), and new methodology such as the approach presented here might improve functional anatomical understanding.

In conclusion, serial intravital imaging of the spleen is a new tool in the investigative toolbox of immunologists that may help shed light on long-lived and dynamic immune reactions in the spleen without perturbing the immune system. It may find use in other areas as well, such as hematology, oncology, and infectious disease research.

## Materials and Methods

### Mice

*Foxp3*^DTR-GFP^ (B6.129(Cg)-*Foxp3^tm3(Hbegf/GFP)Ayr^*/J) breeding pairs were purchased from The Jackson Laboratory (strain no. 016958) (*73*) and then bred in-house. *Foxp3*^DTR-GFP^ mice express a knocked-in human diphtheria toxin receptor (DTR) fused to a green fluorescent protein (GFP) in all Foxp3+ cells without disrupting normal *Foxp3* expression. This allows visualization of T_REG_ and T_FR_ cells. The knocked-in human DTR receptor furthermore allows a targeted ablation of FoxP3+ cells with diphtheria toxin, this approach was however not used in these experiments and the transgenic strain was only used to visualize FoxP3+ cells. To ensure DTR-GFP expression throughout the entire peripheral T_REG_ cell population, female mice were bred homozygously and male mice hemizygously for the X-linked knock-in allele.

Ubiquitin C (UBC) photoactivatable (PA)-GFP (B6.Cg-*Ptprc^a^*Tg(UBC-PA-GFP)1Mnz/J) breeding pairs were purchased from The Jackson Laboratory (strain no. 022486) (*74*) and then bred in-house. UBC PA-GFP mice express photoactivatable GFP under the control of the ubiquitin C promoter. Upon two-photon λ_Ex_ = 830 nm stimulation, PA-GFP switches its absorption maximum from ∼400 to ∼500 nm, yielding a significant intensity increase when excited at λ_Ex_ = 940 nm. UBC PA-GFP mice enable rapid, stable fluorescent labeling of living cells with great microanatomical precision (*74*).

Mice were housed under specific-pathogen-free (SPF) conditions in individually ventilated cages under regulated and stable temperature and humidity. Offspring were weaned at 3 weeks of age and mice were kept on a 12-hour light/dark cycle with standard chow and water ad libitum. All experimental procedures involving animals in this study were approved by local authorities (Animal Experiments Inspectorate, Denmark, license number: 2022-15-0201-01288). Animals were age- (7-14 weeks at initiation of experiments) and sex-matched. Details of experimental and biological replicates are indicated in respective figure legends.

### Resiquimod (R848) treatment

Mice were briefly anesthetized in continuous flow of 4% isoflurane, then treated topically on the right ear with 1 mg R848 (Sigma-Aldrich)/mL acetone. Treatment was performed using a 15 cm, latex- and PVC-free cotton-tipped applicator, which was dipped into the R848 solution until soaked, then rolled on the inner leaflet of the ear. This process was repeated on the outer leaflet of the ear. For intravital observation of autoreactive GC initiation, mice were treated twice before starting the observational period and then treated 3x per week until the end of experiment. For the R848 cessation cohort, mice were treated 3x per week for 4 weeks, then treatment was stopped, and the mice were imaged for two weeks.

### Abdominal imaging window (AIW)

Custom-made winged titanium rings (SOLIDPART Denmark) were adapted (*40*) from the original AIW model (*41*) to specifically enable and simplify serial intravital imaging of abdominal organs using an upright 2-photon microscope. The rings are double rimmed with a diameter of 14 mm excluding the wings. A 0.4 mm deep and 12.2 mm ø wide recess is engraved on top of the ring to fit a glass coverslip (*40*). Before each implantation, titanium rings were cleaned with soap and water, sterilized using 70% ethanol, and then autoclaved individually. Round 12 mm ø wide glass coverslips were sterilized and passivated with oxidation-stable antifouling brushes to ensure biocompatibility of the glass surface while retaining transparency of the implanted AIW over several weeks. The passivation procedure is described below. Glass coverslips were coated several days before window preparation and stored under clean and sealed conditions at −20°C until usage. The day before surgery, using approximately 7 µl of n-Butyl cyanoacrylate glue (Loctite® Super Glue Glass), the coverslip was glued into the engraved recess of the autoclaved titanium ring under laminar airflow, then stored in a sterile-sealed petri dish at room temperature to allow the development of the glue’s full bond strength overnight.

### Glass coverslip passivation

Prior to coating, borosilicate glass coverslips (Menzel Gläser, vwr^TM^, 631-0713) were cleaned by ultrasonication in acetone and 96% ethanol, followed by drying under N_2_ stream. Each side of the coverslips was plasma cleaned for 15 min using a Diener Femto Plasma Etcher at 100 W at ≈100 mTorr, and an O_2_ flow of 30 SCCM was used to activate the surface for attachment of covalent silane linkers. The plasma-cleaned coverslips were then immediately transferred to a sterile Petri dish in a sterile environment under a laminar airflow bench, and were transferred to a sterile 50 mL Falcon tube containing 0.2 μm filtered solution of 0.1 mg/mL Poly(acryl-amide)-g-(PMOXA, 1,6-hexanediamine, 3-aminopropyldimethylethoxysilane)) PAcrAm-g-PMOXA (NH2,Si), (SuSoS AG - Switzerland) dissolved in 1 mM HEPES buffer at pH 7.4. According to an adapted protocol from Ogaki et al. (*75*), the tubes were incubated for 24 h in a water bath at 80°C to achieve ultra-dense coating and optimize the longevity of the antifouling coating. After 24 h of incubation, the coverslips were carefully removed from the solution, rinsed with an excess of 0.2 μm filtered deionized H_2_O, and dried under N_2_ stream. The low-temperature PMOXA coating was prepared similarly but by incubation at room temperature.

### *In vitro* passivation comparison and analysis

RAW 264.7 macrophages (passages 4-12) and NIH 3T3 fibroblasts (passages 11-18) were used to compare the cellular attachment to the passivated coverslips. The cell cultures were maintained in Dulbecco’s modified Eagle’s medium (high glucose DMEM with GlutaMAX, Thermo Fisher Scientific). Cells were incubated at 37°C with 5% CO_2_ and supplemented with 10% (v/v) fetal bovine serum (FBS), 100 mg/ml penicillin, and 100 mg/ml streptomycin. Cells were passaged by trypsinization when approaching 80% confluence.

The passivated glass coverslips were prepared as described above and were attached to Greiner bio-one bottomless 96-well plates (ID: 655000) using laser-cut double-sided adhesive (ARcare 90106NB) as previously described (*76*).

The cell experiments were performed in the same medium, including 10% FBS, which was used for culturing at a seeding density of 3000 cells/cm^2^. Each day, 2/3 of the medium in each well was replaced with fresh medium. On days 4 and 8, an additional 3000 cells/cm^2^ were seeded to ensure the supply of fresh viable cells on the non-adhering surfaces.

After 14 days, the cells were fixed for 10 minutes in 4% PFA and permeabilized in 0.2% Triton-X 100 for 10 minutes before staining for nuclei with 300 nM DAPI (Sigma-Aldrich) and for F-actin with 50 nM Atto-488 Phalloidin (AttoTec, Germany) for 30 minutes. The central 50% of each well was imaged using the high-content imaging system ImageXpress Pico (Molecular Devices) with a 20x objective. The cell experiments were repeated independently three times, with each independent repeat having two experimental repeats per condition. Nuclei images were segmented using Cellpose v3.0 (*77*). Pretrained nuclei model on a GPU (Tesla T4) enabled on Google Colab notebook using a custom-made Python script to count the number of the adherent cells. Binary thresholding of pixel intensities above the background gray value of F-actin images was used to estimate the percentage of the surface covered by the cells.

### AIW implantation over the spleen

A surgical protocol was adapted from the AIW implantation over the left kidney (*39*). Surgical instruments were sterilized in a glass bead sterilizer and all surgical surfaces including the heating plate were cleaned using 70% ethanol. Additionally, a sterile, disposable surgical drape was used to cover the heating plate. Mice were anesthetized with isoflurane in medical air and 50% oxygen (induction: 3.5% isoflurane; maintenance 1.2–1.8% isoflurane; flow rate: 0.6 – 1.2 L/min, Anesthetic Vaporizer UNO300VAP). Mice received analgesia 15 min prior to surgery with Buprenorphin in 0.9% NaCl (0.1 mg/kg bodyweight i.p.) and were placed on the draped and cleaned heating plate. A nose cone was used to maintain anesthesia, and protective eye ointment (Visc-ophtal øjengel 2mg/g, Orifarm) was used to prevent corneal ulceration. The surgical area on the left flank including surrounding fur was disinfected with chlorhexidine (local hair removal the day prior to surgery). The AIW was placed between the left ribcage and left hind leg, ventral to the spine (Fig. S4). After loss of pain reflexes, a 1 cm long dorsoventral incision in the left flank was made, cutting under vision through skin, fat tissue and muscle layer, respectively (Fig. S4A-C). Minor bleeding was stopped with a hemostatic dental sponge (Spongostan, Ethicon). Starting from dorsal, a purse-string suture was set around the incision using a 6-0 non-resorbable monofilament prolene suture (Ethicon), connecting muscle and skin layer, while avoiding inclusion of fat tissue (Fig. S4D). Then the spleen was carefully mobilized from the abdominal cavity by rolling gently with a sterile cotton swab over the perisplenic fat, combined with a soft pull on the perisplenic fat using dull tweezers. The surgical field was regularly irrigated with droplets of sterile 0.9% NaCl to reduce friction during spleen mobilization (Fig. S4E). After successful spleen mobilization, skin around the incision was covered in sterile packed non-woven cotton tissue strips pre-wetted with sterile 0.9% NaCl (Fig. S4F). Then 10 µl of n-Butyl cyanoacrylate glue (Loctite® Super Glue Glass), was applied in the inner borders of the AIW sink (where glass coverslip meets the titanium ring) (Fig. S4G). The AIW was then glued onto the spleen in one smooth movement and held still for 5-7 min to allow polymerization of the glue (Fig. S4H). Finally, the skin was carefully inserted into the grove between the two rims of the implant and the purse-suture was tightened and secured with surgical knots (Fig. S4I+J).

### Sham surgery

Sham-operated mice were handled similarly to AIW operated mice. After the incision, the spleen was immediately mobilized and exposed similarly to AIW mice without laying a purse string suture beforehand. After the spleen was guided back into the abdominal cavity, muscle and skin layer were closed after another with approximately 10 single stitches per layer. A 6-0 resorbable vicryl coated suture (Ethicon) was used for the muscle layer and a 6-0 non-resorbable monofilament prolene suture (Ethicon) for the skin layer.

### Pre- and postoperative management

To acclimatize mice to the taste of analgesic drinking water, important to avoid dehydration immediately after surgery, normal drinking water was exchanged with buprenorphine (0.009 mg/ml) drinking water already two days *before* surgery. Chow was additionally soaked with buprenorphine drinking water to enhance adaptation to taste. One day before surgery, the hair in surgical field (ca. 2 cm x 2 cm) on the left flank was removed under anesthesia. First with an electrical shaver, then with depilation cream. A thin layer of moisturizing skin cream was applied to soothe the skin after hair removal.

Immediately after surgery, mice received meloxicam (1 mg/kg bodyweight s.c.) for additional pain-relief and anti-inflammation. Postoperative analgesia (0.009 mg/ml buprenorphine drinking water and 1 mg/kg bodyweight meloxicam s.c./day) was implemented for 2 additional days after the surgery day. Immediately after surgery, mice were placed in a fresh and clean cage under an infrared heating lamp for 30 - 60 min. recovery. Mice were single-housed and monitored daily for appropriate wound healing and general thriving (e.g. shiny fur, active behavior, grooming tendencies, etc.). Moisturizing skin care was carried out whenever deemed necessary. Mice were weighed daily (SCOUT™ STX2201, High-Performance Portable Balance) to monitor their recovery. To support a swift and pain-free recovery, mice were provided with soaked chow (buprenorphine drinking water) in the first 3 post-operative days and easily accessible dry chow on the cage bottom throughout the entire imaging period. The cages were equipped with modified enrichment to avoid mice getting stuck with the wings of the implanted AIW after surgery: Tubes for tunnel handling with bigger diameter than regular paper rolls. The entry of paper houses was cut wider. Long entangling nesting material was replaced with soft cotton buds and soft <1.5 cm short bedding material. Chewing sticks/balls were used for further enrichment. Mice were given 1-3 days to recover after surgery before the first imaging session.

### Intravital labeling techniques and intravital cell identification

To label FDCs and thereby mark the light zone of GCs, mice were injected with 2 µg αCD35-iFluor647 in 100 µl sterile PBS i.v. 24 h before each imaging time point. αCD35 (clone 8C12, BD Biosciences 558768), was conjugated to iFluor647 (ABD-1031, Nordic Biosite) using an in-house labeling protocol (final concentration 0.43 mg/ml and labeling density 12). T_REG_ cells were visualized based on GFP expression of the fused DTR in FoxP3+ cells (*73*). T_FR_ cells were furthermore identified based on their localization in GCs (*50*). This included GFP+ cells localized within the FDC-labeled area, as well as areas in very close proximity, especially those close to TBMs. Different macrophage populations, such as TBMs, were easily identified based on bright vacuolar autofluorescence and their location. Further microanatomical landmarks of follicles, such as perivascular-T (PT)-tracks (*58*), were visualized by large collagen bundles close to follicles, detected by second harmonic generation (SHG) imaging (*78, 79*). The marginal zone was labeled only for fresh spleen explant imaging and was visualized by injecting 2 µg CD169-PE (142404; Biolegend) diluted in 100 µl PBS i.v. 10 min before euthanasia.

### Serial intravital 2-photon microscopy

For intravital imaging, mice were anesthetized with isoflurane in medical air enriched with 33% oxygen (induction: 3.5% isoflurane; maintenance 0.7 –1.5% isoflurane; flow rate: 0.6 – 1.2 L/min, using either SomnoSuite apparatus (Kent Scientific, United States) or Anesthetic Vaporizer UNO300VAP). After induction, mice received fluid therapy, 100 µl 0.9% NaCl s.c. and eyes were protected with eye ointment (Visc-ophtal øjengel 2mg/g, Orifarm). The AIW was wiped with 70% ethanol and lens paper wrapped around a cotton swab. Then the mouse was mounted in a custom 3D-printed holder for upright intravital imaging (comparable as described in (*40*)) and placed on a servo-controlled heating plate. The focus was then found using epifluorescent light.

Microscopy was carried out with an Olympus FVMPE-RS multiphoton system microscope with Fluoview FV31S software, equipped with an integrated x25 water immersion objective with an isolated tip and numerical aperture of 1.05 (Olympus, XLPLN25xWMP2: WD 2.00 mm), a MaiTai DeepSee Olympus laser (Spectra Physics), specifically tailored for the FVMPE-RS system and 2 high-performance multi-alkali PMT and 2 GaAsP PMT detectors. Images were acquired with two sequential excitation wavelengths, λ _Ex_ = 840 nm and λ _Ex_ = 940 nm, to capture distinct spectral features: λ _Ex_ = 840 nm was used to excite iFlour647 (CD35) and λ _Ex_ = 940 nm was used to excite GFP (FoxP3) and observe SHG (collagen). Fluorescent emission was collected in 4 channels (to balance signal to autofluorescence) in both excitation tracks using the following filter sets: Ch 1 & 2: 650/60 & 573/75, separated by Olympus FV30-SDM570 primary emission beam splitter from Ch 3 & 4 (GaAsP): 520/30 & 452/45. An IR short pass filter, Olympus BA685RXD, was used to only allow emission light below 685 nm.

Individual GCs were acquired in volumetric image stacks with 5 µm step size between imaging frames, reaching a total depth of up to 220 µm – measuring from capsule to deepest imaging point. At λ_Ex_ = 840 nm, images were acquired in galvo scanning mode without averaging using 2 µs dwell time/pixel in 800 x 800 pixels resolution and at λ_Ex_ = 940 nm with 3 x line averaging using 4 µs dwell time per pixel in 800 x 800 pixels resolution. Overview maps of the spleen area within the window’s field of view were manually outlined at around 120 µm below the capsule to set an imaging grid using the Multi-area Time Lapse (MATL) function in Fluoview to finally generate a 3D stitched overview of the entire imaging window during the ongoing imaging session. Single tiles of the 3D map were acquired in the MATL function at λ_Ex_ = 840 nm without averaging using 2 µs dwell time per pixel in 512 x 512 pixels resolution with 50 µm step size from 0 µm to 250 µm (0 µm marking the capsule) and 10% overlap of tiles. Detector Gains were adjusted in the beginning of each serial imaging cohort to balance fluorescent emission of interest and autofluorescence and then kept unchanged during each experiment. Laser Power was increased exponentially in depth to detect all fluorophores of interest throughout the entire 3D stack, while avoiding signal saturation, phototoxicity and bleaching.

Each intravital 2-photon imaging session lasted around 90 – 120 min per mouse. Appropriate respiration rate in anesthesia during image acquisition was monitored using an infrared camera (SVPRO 1080P Night Vision USB Camera CMOS OV2710 IR LED Infrared Webcam), and protective eye ointment was reapplied approximately every 30 min. After each imaging session, mice were rehydrated with another 100 µl 0.9% NaCl s.c. and provided with soaked chow. Mice usually woke up and started grooming themselves 2 - 5 min after anesthesia was withdrawn. Mice were imaged at 6 time points over 16 days.

### Imaging analyses of intravital images

To compare volumetric changes in GCs in the spleen over time reliably, 3D imaging stacks from all six imaging days of individual GCs were standardized in stack position and stack size using Fiji (v2.14.0/154f) (*54*). To standardize the imaging stacks, landmarks of capsule, vasculature and splenic trabeculae were manually matched to the same frame level. Based on matched frame positions, the entire 3D stack was reduced to the largest Δframes that were found reproducibly throughout all days. Standardizing imaging stacks also served as a checkpoint to reconfirm that the same GCs were observed over time. Each longitudinal imaging set was finally visualized with the Multi Stack Montage plugin from the PTBIOP Update Site (*80*) to ensure consistency in stack standardization. Synchronization of windows is also possible with the built-in Fiji plugin “SyncWindows” (*54*).

To visualize and analyze GCs in depth of the spleen, the imaging data was displayed in an orthogonal view using Reslice function, followed by Z-projection in Fiji (*54*).

To generate a quick, but reliable overview of the FDC-network size at respective imaging time points, an outline of the FDC-network was manually drawn and measured (area) on an *Average Z-Projection*, built-in Z function in Fiji (*54*) of the λ _Ex_ = 840 nm track. Only frames with positive CD35 staining were selected for Z-projection to avoid a distracting overlay of red pulp macrophages above the FDC network and facilitate visualization (Fig. S1). Before using the freehand tool to outline the FDC network, salt and pepper noise was reduced by applying the “Despeckle” function in Fiji (*54*). Manual outline was eased by activating channel 1 (CD35+ and macrophages) and channel 3, while applying complementary pseudo colors (magenta and green) to both channels. Finally, the measured area was normalized by the measurement of the first observation timepoint.

Image preprocessing for T_FR_ cell visualization was carried out entirely in Fiji and no additional plugins were required (*54*). For T_FR_ cell visualization, both excitation tracks were merged to eventually identify FoxP3+ (GFP+) cells in GCs. GC were identified based on their light zone with FDC labeling and the entire GC area was estimated based on TBM localization closely around the FDC network. Before merging channel 1 from λ _Ex_ = 840 nm track and channel 2-4 from λ _Ex_ = 940 nm track, background illumination from the respective channels was individually corrected by rolling ball background subtraction (rolling ball radius 50 pixels). Then, salt and pepper noise of the newly merged image was reduced by applying the “Despeckle” filter. Lastly, the “Maximum 3D” filter was applied to further enhance the GFP-signal of single FoxP3+ cells (T_REG_ and T_FR_ cells) for better visualization.

All representative images underwent image processing as described in respective figure legends.

### Tissue Harvest

For tissue harvest, mice were anesthetized with isoflurane and euthanized by decapitation. First, blood was collected for serological and flow cytometric analyses. For serum preparation, blood coagulation was allowed for 30 min at room temperature, then supernatant was collected after centrifugation at 3,000 *g* for 10 minutes followed by 20,000 *g* for 3 minutes. Serum samples were stored at −80°C until analysis. Blood for flow cytometry was collected in 200 µl PBS with 5 mM EDTA and stored on wet ice until tissue processing. IngLN were removed and cleaned from surrounding fat tissue. Left (ipsilateral) and right (contralateral) IngLN from mice that underwent any kind of surgery, were kept separate. IngLNs were weighed on a precision scale (Mettler Toledo MS303S) and then either fixed in 4% w/v paraformaldehyde (PFA) for histological analyses or stored in FACS buffer (PBS, 2% heat-inactivated fetal calf serum (FCS), 1 mM ethylenediaminetetraacetic acid (EDTA)) on wet ice until tissue processing for flow cytometry. After 24 h of fixation, IngLN for histological analysis were transferred into 30% sucrose in PBS containing 0.1% sodium azide, incubated overnight, then thoroughly and gently cleaned with fine tweezers under a light microscope and embedded in OCT and frozen at −80°C until analysis. Spleens were carefully removed and either freshly prepared for immediate 2-photon imaging or stored in ice-cold FACS buffer for flow cytometry. Spleens from AIW mice were taken out with attention to not damaging the parenchyma while separating the spleen from the glass coverslip. AIW-spleens were further divided into parts attached to the window (and subject to serial imaging) and parts that were distal to the AIW.

### Flow Cytometry

For flow cytometry, blood, spleen and IngLNs were processed following standard protocols: ca. 4 mm thick coronal spleen sections or IngLNs were mechanically macerated in 500 µl ice-cold FACS buffer by reusable pestles. Cell suspensions were then filtered through 100 µm cell strainers. Following filtration, spleen samples were centrifuged at 200 *g* at 4°C for 5 min. After supernatant removal, cells were resuspended in 400 µl red blood cell (RBC) lysis buffer (155 mM NH_4_Cl, 12 mM NaHCO_3_, and 0,1 mM EDTA) at room temperature for 5 min. The reaction was stopped with 1000 µl FACS buffer, then the suspension was centrifuged again at 200 *g* at 4°C for 5 min, the supernatant was removed, and the cells were resuspended in FACS buffer. The blood samples were underlayered with 1 mL of Lympholyte M (CedarLane Labs) and centrifuged 25 minutes at 600 *g* at room temperature. The peripheral blood mononuclear cell (PBMC) layer was aspirated with a pipette and resuspended in 1 ml FACS buffer, then centrifuged at 200 *g* at 4°C for 5-10 minutes. The supernatant was removed, and the pellet resuspended in FACS buffer. All cell suspensions were stored on ice until plated.

Before adding 100 µl of cell suspension to wells, 20 µl of 1:50 diluted Fc-block (2.4G2), was added into wells of a 96-well plate. Antibodies and a viability dye were diluted in a 50:50 mix of FACS buffer and Brilliant Stain Buffer (BD Horizon™) and 100 µl of the antibody mix was added to each plated sample and incubated for 30 min on ice. Then the plate was centrifuged at 200 *g* for 5 min 4°C, the supernatant was removed, and the pellet resuspended in 200 µl FACS buffer. The last step was repeated, and the samples were then freshly analyzed using either a BD LSRFortessa^TM^ Cell Analyzer equipped with 4 lasers (405 nm, 488 nm, 561 nm, 640 nm) and 16 fluorescence detectors (BD Biosciences, San Jose, CA) or a Novocyte Quanteon 4025 equipped with 4 lasers (405 nm, 488 nm, 561 nm and 637 nm) and 25 fluorescence detectors (Agilent, Santa Clara, CA). Data were acquired in either BD FACSDiva Software version 8.0.2 (LSRFortessa, BD Biosciences, San Jose, CA) or NovoExpress version 1.6.2 (Quanteon, Agilent, Santa Clara, CA) and analyzed in FlowJo^TM^ version 10.8.1 software (BD Lifesciences). Gating strategies can be found in Fig. S3. Antibodies including dilutions used for flow cytometry experiments can be found in table S2.

### Vibratome sections and 2-photon imaging of fresh spleen explants

Spleens for 2-photon explant imaging were placed on ice and processed immediately after harvest. Using a vibrating blade microtome (Leica VT1200 S), 400 - 800 µm thick transversal spleen slices were prepared. Beforehand, the vibratome tray was filled with wet ice and an object glass was mounted on the specimen plate with reusable sticky tack and leveled with a spirit level. Spleens were then briefly blot-dried on filter paper and then superglued on the leveled object glass. Cutting parameters: speed: 0.18 mm/s; amplitude: 1 mm. Both spleen and blade were irrigated with PBS droplets during the entire cutting process and spleen slices were prevented from folding using a fine brush. Slices were mounted in PBS between two coverslips to allow imaging from both sides, and vacuum grease was used to seal the chamber. Slices were imaged immediately at most times, could however also be stored in the dark at 4°C overnight without an appreciable drop in imaging quality the next day.

Fresh explant 2-photon microscopy was carried out on the same multiphoton system with similar hardware settings as during intravital microscopy. Images were acquired with two sequential excitation wavelengths, λ_Ex_ 840 nm and λ_Ex_ 940 nm, to capture distinct spectral features: λ_Ex_ 840 nm was used to excite iFlour647 and PE and λ_Ex_ 940 nm was used to excite GFP, PE and observe SHG. 3D image stacks were acquired using the MATL tile scan function in Fluoview (2×2 - 4×4 single tiles) to capture entire white pulp areas. Stack acquisition with 5 µm step size reaching a total depth of up to 200 µm (measuring from cut surface to deepest imaging point). All images were acquired in galvo scanning mode with 3 x line averaging, 4 µs dwell time/pixel at 800 x 800 pixels resolution. Detector Gains were adjusted in the beginning of each experimental cohort to balance fluorescent emission of interest and autofluorescence. Laser Power was exponentially increased in depth and adapted to avoid signal saturation and bleaching while still illuminating all fluorophores of interest throughout the entire 3D stack. Laser Power intensity was set lower than during comparable intravital scans.

### Ki67 staining and 2-photon imaging of stained vibratome sections

To visualize the biological dark zone of each imaged GC of the fresh spleen explants, thick slices were stained with Ki67-AlexaFluor488 (monoclonal (SolA15), eBioscience™) subsequently to imaging. After imaging, slice orientation and approximate localization of imaged area were photo documented. Then, spleen slices were carefully removed from the imaging chamber and placed in an Eppendorf tube with 4% PFA overnight at 4°C. After fixation, slices were transferred into a tube with PBS and 0.1% sodium azide at 4°C until further permeabilization and staining. The slice that remained glued to the objective glass was occasionally imaged as well, and a fixation, storage and staining protocol for thick spleen sections glued to an objective glass was developed using 50 mL tubes, vacuum grease and custom 3D-printed humidified staining chambers. Before staining, spleen slices were washed in PBS (3 x 15 min) on a shaking table at room temperature in the dark. Then samples were blocked in PBS containing 5% FBS and 0.5% Triton X-100 overnight on a shaking table at room temperature. The following day, blocking buffer was exchanged with staining mix, Ki67-AlexaFluor488 (clone: SolA15) 1:70 in PBS containing 2.5% FBS and 0.5% Triton X-100. After 48 hrs incubation on a shaking table at room temperature, samples were washed in PBS (3 x 15 min), and each slice was mounted between two coverslips and vacuum grease in the same position as the fresh explant (with the help of the previous photo documentation to facilitate re-finding the same imaged areas). *FoxP3*-GFP signal disappeared in the staining process and was only visible in fresh spleen explants. Intravital labeling of αCD35-iFluor647 and αCD169-PE largely withstood the fixation and staining process and supported reidentification of imaged areas from fres h explants. Reidentification of the same areas was usually already achieved through oculars by reidentification of different CD169-PE staining patterns excited by epifluorescent light. Exact location was then confirmed with 2-photon excitation. To facilitate reidentification of previously imaged areas, laser marks can additionally be set in fresh explants and re-found in stained slices (Fig. S5A-C) as also described by Ritsma et al. (*81*). Laser marks were set using λ_Ex_ 840 nm, 100 µs dwell time/pixel for 500 ms long per mark. The fragile consistency of fresh spleen tissue prevented a safe preservation of all set laser marks and hence multiple marks had to be set. The tissue shrank slightly during the fixation and staining process (Fig. 8F-I).

Images were acquired in λ_Ex_ 840 nm excitation, to capture especially AlexaFluor488, but also iFlour647, PE and distinct collagen fibers. 3D image stacks were acquired using the MATL tile scan function in Fluoview in galvo scanning mode with 3-line averaging using 4 µs dwell time/pixel at 512 x 512 or 800 x 800 pixels resolution and stitched post image acquisition. Detector Gains were adjusted in the beginning of each experimental cohort to balance fluorescent emission of interest and autofluorescence. Laser Power was adjusted to avoid signal saturation and bleaching while still illuminating all fluorophores of interest throughout the entire 3D volume. Laser Power was increased exponentially in depth, but intensity was lower than during comparable intravital scans.

All representative images underwent image processing as described in figure legends. Additionally, representative images from explanted spleens both λ _Ex_ = 840 nm and λ _Ex_ = 940 nm, were registered using the BigWarp plugin (*82*) in Fiji for manual landmark-based image alignment. Autofluorescent macrophages (red pulp macrophages and TBMs) served as landmarks.

### Confocal microscopy of inguinal lymph nodes

A Cryostar NX70 Cryostat (ThermoFisher) was used to cut 20 µm thick IngLN sections from the middle of each LN. Sections were mounted on SuperFrost+ glass slides (Epredia, REF J1800AMNZ). IngLN sections were acetone fixed (Merck, catalogue number 1000141000). Briefly, the spleen samples were rinsed in PBS and fixed in acetone for 10 minutes at room temperature (RT). Thereafter slides were washed two times with PBS 0.1% w/v sodium azide. Biotinylated antibodies were diluted in staining buffer (PBS, 2% v/v FBS, 0.1% w/v sodium azide and spun for 10 min at 10,000 *g* at 4℃ to avoid antibody aggregation. Slides were stained for 4 hours at 4 ℃ in darkness. Slides were washed once with staining buffer for 5 minutes and washed three times for 5 minutes with PBS 0,01% v/v Tween-20 (Merck, product no.: 8.17072). Then, primary conjugated antibodies and primary conjugated streptavidin were diluted in staining buffer (PBS, 2% v/v FBS, 0.1% w/v sodium azide) and spun for 10 min at 10,000 *g* at 4 ℃ to avoid aggregation. Slides were then finally stained overnight at 4℃ in darkness. Slides were washed once with staining buffer, and three times for 5 minutes with PBS 0,01% v/v Tween-20 spot-dried and mounted using Fluorescence Mounting Medium (S3023, Dako). After drying, slides were sealed airtight using nail polish and stored in darkness at 4℃ until imaged. All imaging was done using a Zeiss LSM800 confocal microscope with a 10x objective, equipped with Airyscan. The following antibodies were used: CD35-iFlour647 (1/500 – in-house conjugation clone 8C12, BD Biosciences 558768 and ABD-1031, Nordic Biosite), CD169-PE (1/500 – (142404; Biolegend)), and Thermo Fisher Scientific, S32354), Lyve-1-eFlour450 (1/300 – Invitrogen, clone: ALY7, 48-0443-82). Confocal images were processed and analyzed using Fiji (v2.14.0/154f) (*54*). Area measurement of the entire LN was carried out by a manual outline. All LN-lobes were included and extra nodular lymphatic vasculature and fat tissue excluded. The area of the intranodal lymphatic vasculature was measured by the Lyve-1 staining area after clearing the outside of the manual LN outline. Auto-thresholding on the Lyve-1 staining in default mode, was used to create a binary mask. The noise of the mask was reduced by applying the Gray Morphology Filter (radius=1, type=circle, operator=erode) and median filter (Despeckle). The area-coverage of the mask was measured.

### Statistical Analyses

Mice in different groups were randomly assigned, and studies were non-blinded by nature. A few animals and imaging data sets were excluded due to technical obstacles incl. bleeding and extreme tissue remodeling, respectively. Outliers were not excluded, and there was no statistical pre-determination of sample size. Statistical analyses were performed using GraphPad Prism (v. 10.1.1). QQ plots were used to determine degree of normal distribution, the need for data transformation, and to choose a suitable statistical methodology. Statistical analysis was performed on raw data or on log-transformed data when QQ-plot indicated log-normal distribution. Statistical significance was determined using two-tailed paired t-tests, and in case of grouped analyses, parametric or non-parametric tests depending on normality and homogeneity of variances (one-way ANOVA, two-way ANOVA or Kruskal-Wallis’s test). Data are expressed as means ± SD. P values < 0.05 were considered as significant and only p values < 0.05 are shown in graphs unless indicated otherwise. Test indicated in respective figure legend with n and p-values.

## Acknowledgements

We thank the Laboratory Animal Facility, the Bioimaging Core Facility, and the FACS Core at Aarhus University for excellent technical assistance and support. We thank the technicians at the Department for Biomedicine for excellent support in custom 3D printing.

## Author contributions

Conceptualization: LP, TRW, SED.

Methodology: LP, TRW, AS, KSK, CFH, DoS, IMS, AP.

Investigation: LP, TRW, AS, KK, CFH.

Visualization: LP, AS, SED.

Supervision: DuS, SFG, SED.

Writing—original draft: LP, AS, SED.

Writing—review & editing: All authors.

## Data and materials availability

Data and code will be made available by the corresponding author upon request.

## Supplementary material

Figure S1-5

Table S1+2

---

**Supplementary Figure 1.**
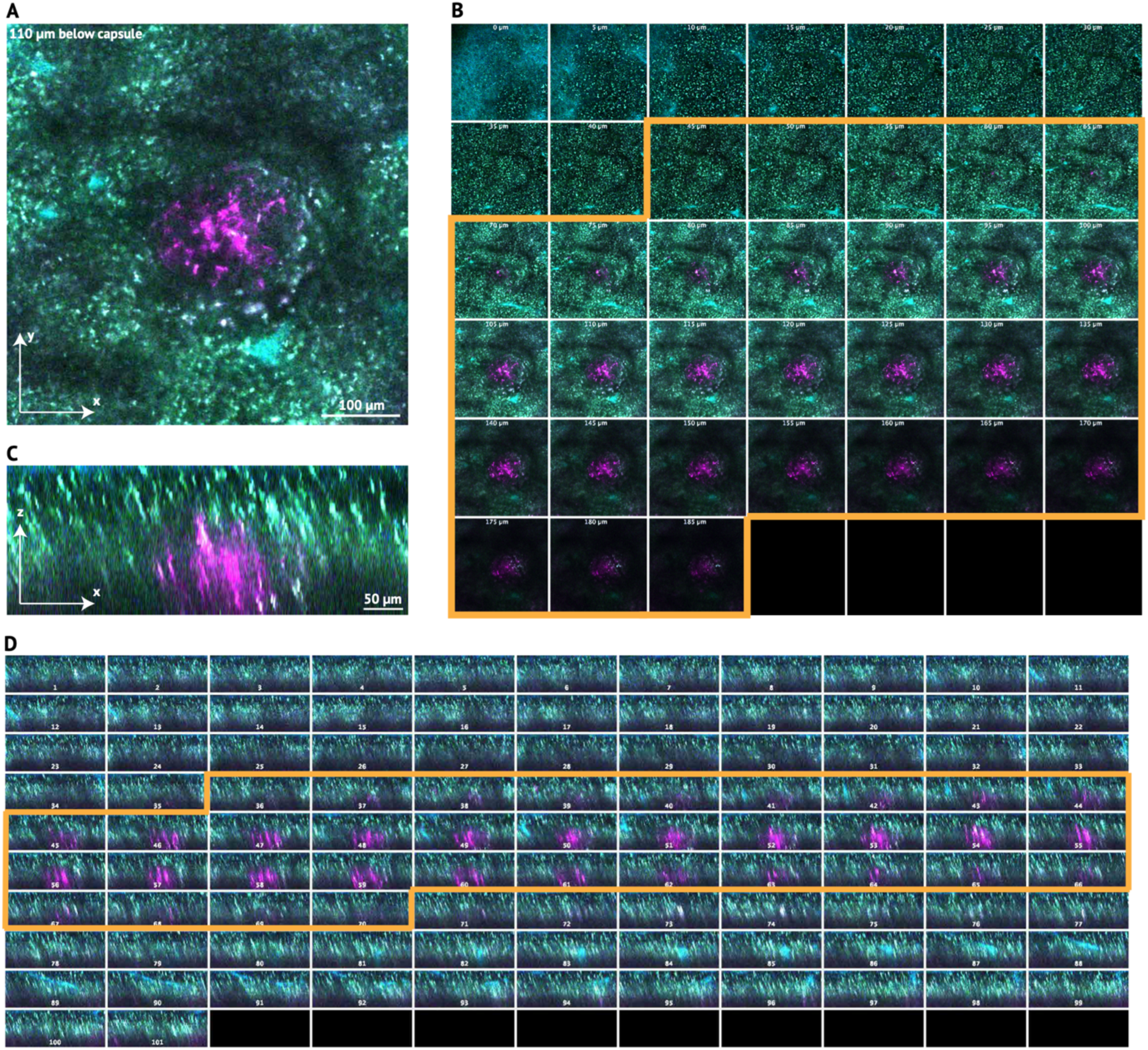
Visualization of 3D datasets of the spleen. **(A-D)** Different displays of a 3D imaging stack to visualize 3D remodeling of the same GC using Fiji. **(A)** Single Z-plane (image frame) of a 3D image stack. Only X and Y dimensions are visible. Scalebar: 100 µm. (**B)** Montage of all single Z-frames of the XY oriented 3D stack (from “A”), starting from capsule, 5 µm step size, until 185 µm below capsule. Orange outlines show what frames were manually selected for 2D Z-projections used for FDC network or GC remodeling analysis. Only frames with staining/areas of interest were selected to be included in the Z-projection to avoid overlay of autofluorescent macrophages, which would complicate manual image analysis. **(C)** Single plane (image frame) of the same 3D image stack as in “A”, orthogonally resliced. Only Z and Y dimensions are visible. Scalebar: 50 µm. **(D)** Montage of all single frames of the ZY oriented 3D stack (from C). Similar principle of the selected frames, outlined in orange (as in B).

**Supplementary Figure 2.**
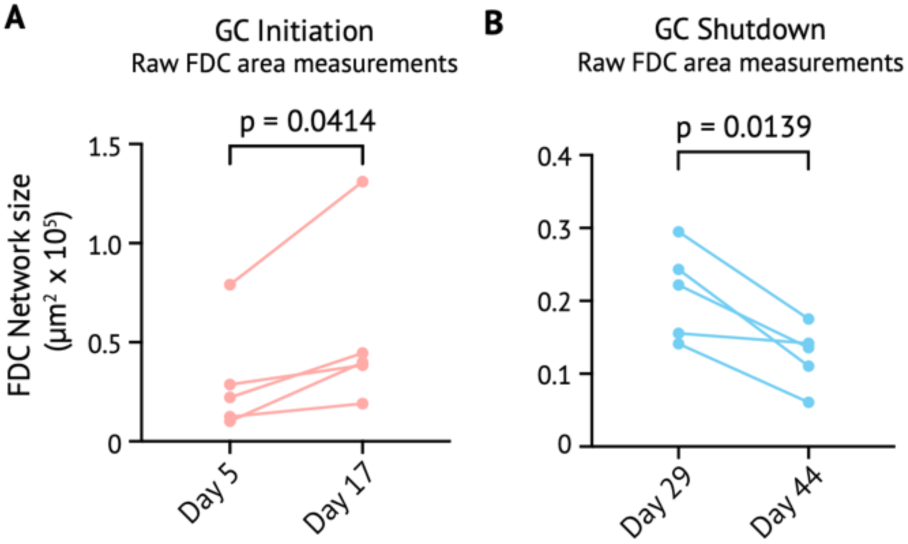
FDC Remodeling. **(A+B)** Raw data of measured FDC area of a 2D Z-projection of imaging frames that were positive for CD35 labeling (compare with Fig. S1C). Raw data is not standardized to the first observation day as in Fig. 6C+D. Usually, measured area is normalized to day 5 - initiation (A) and day 29 – shutdown (B), respectively, set as 100% starting point.

**Supplementary Figure 3.**
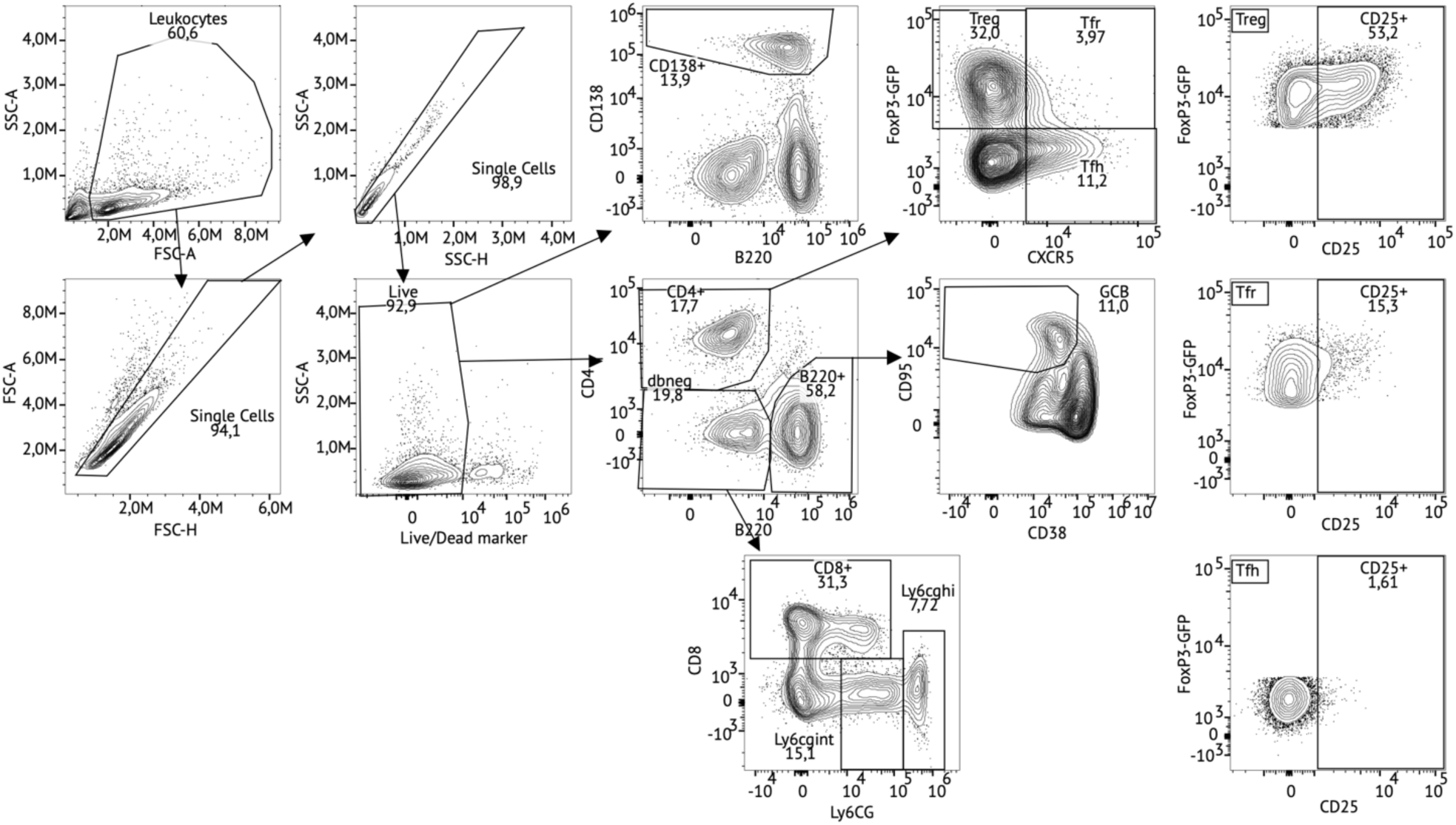
Flow Cytometry - Gating strategy. Gating strategy of flow cytometry data, applied on spleen and IngLN datasets. Leukocytes were gated based on FSC-A and SSC-A, then singlet gating was applied using FSC height versus area and SSC height versus area, followed by live gating based on live/dead marker exclusion. From live, singlets, plasmablasts and plasma cells were gated as CD138. Similarly, from live, singlets, CD4+ T cells, B220+ B cells, and double negatives were gated. CD8+ cells, Ly6cg intermediate and high populations were sub-gated from the double negative gate. GCB cells were sub-gated from the B220+ B cell population. Finally, CD4+ cells were gated as FoxP3+CXCR5-Tregs, FoxP3+CXCR5+ Tfr, and FoxP3-CXCR5+ Tfh. Each of the CD4+ sub-populations were assessed for CD25 positivity.

**Supplementary Figure 4.**
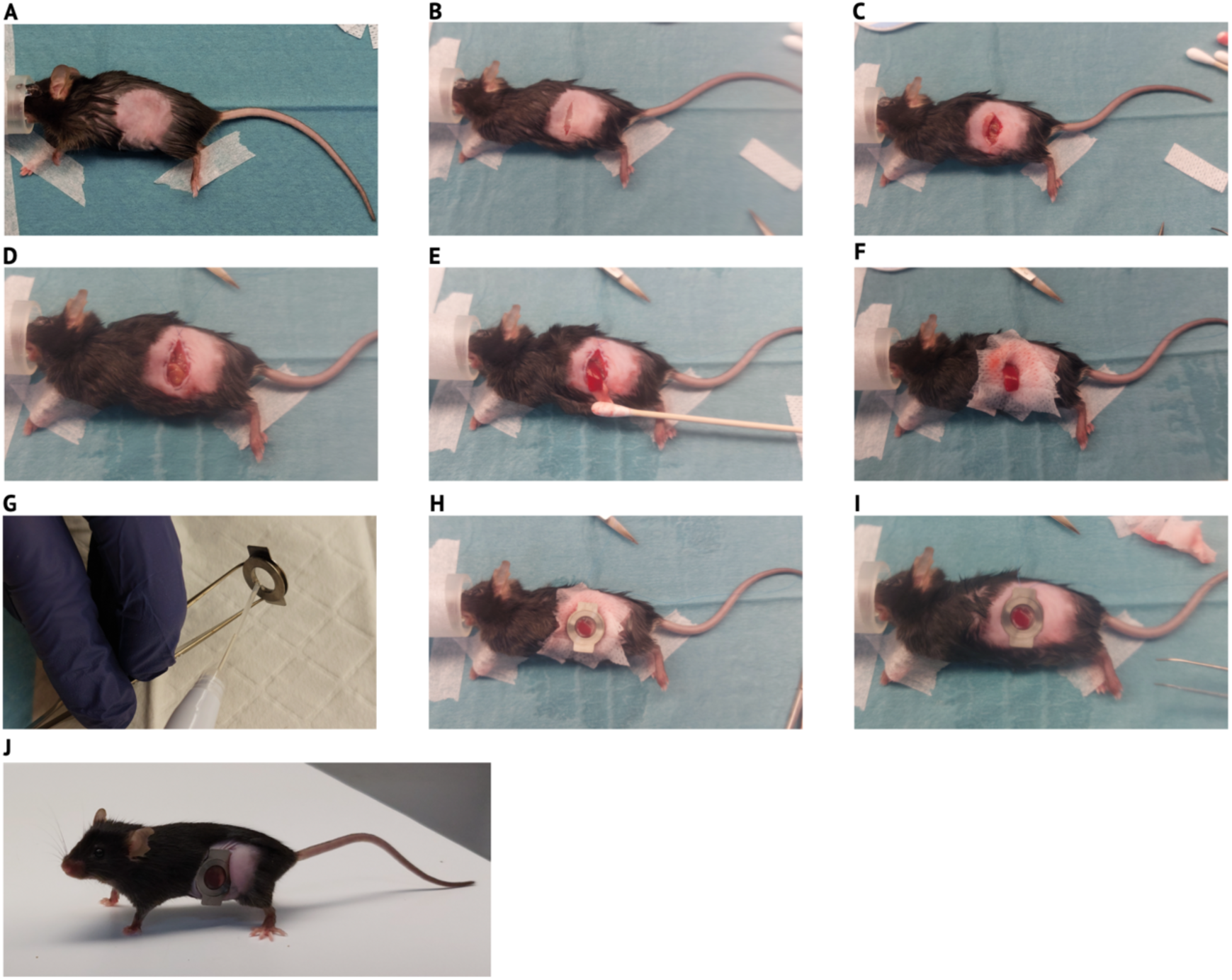
Surgical implantation of an abdominal imaging window (AIW) over the spleen. **(A)** Hair removal and skin disinfection. (**B)** Skin incision. (**C)** Incision of fat and muscle layer. (**D)** Setting purse string suture. (**E)** Gentle and careful spleen mobilization. (**F)** Skin protection with cotton tissue (to avoid glue touching the skin). **(G)** Adding glue on inner window border. (**H)** Placement of AIW with glue onto spleen. (**I)** Inserting skin-muscle layer into the rim of the window, followed by tightening and knotting of suture. (**J)** Awake mouse approx. 1 week after surgery.

**Supplementary Figure 5.**
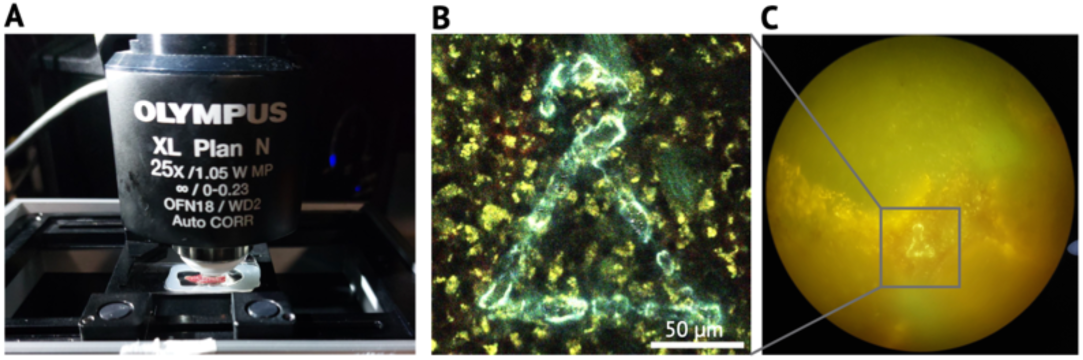
Laser marks in spleen explants for area re-identification after staining. **(A)** Fresh thick spleen section under 2-photon microscope. (**B)** Laser mark set in fresh spleen explants. Laser mark set and image acquired with 2-photon microscope. Scale bar: 50 µm. (**C)** Laser mark identified through the ocular.

**Supplementary Table 1.**
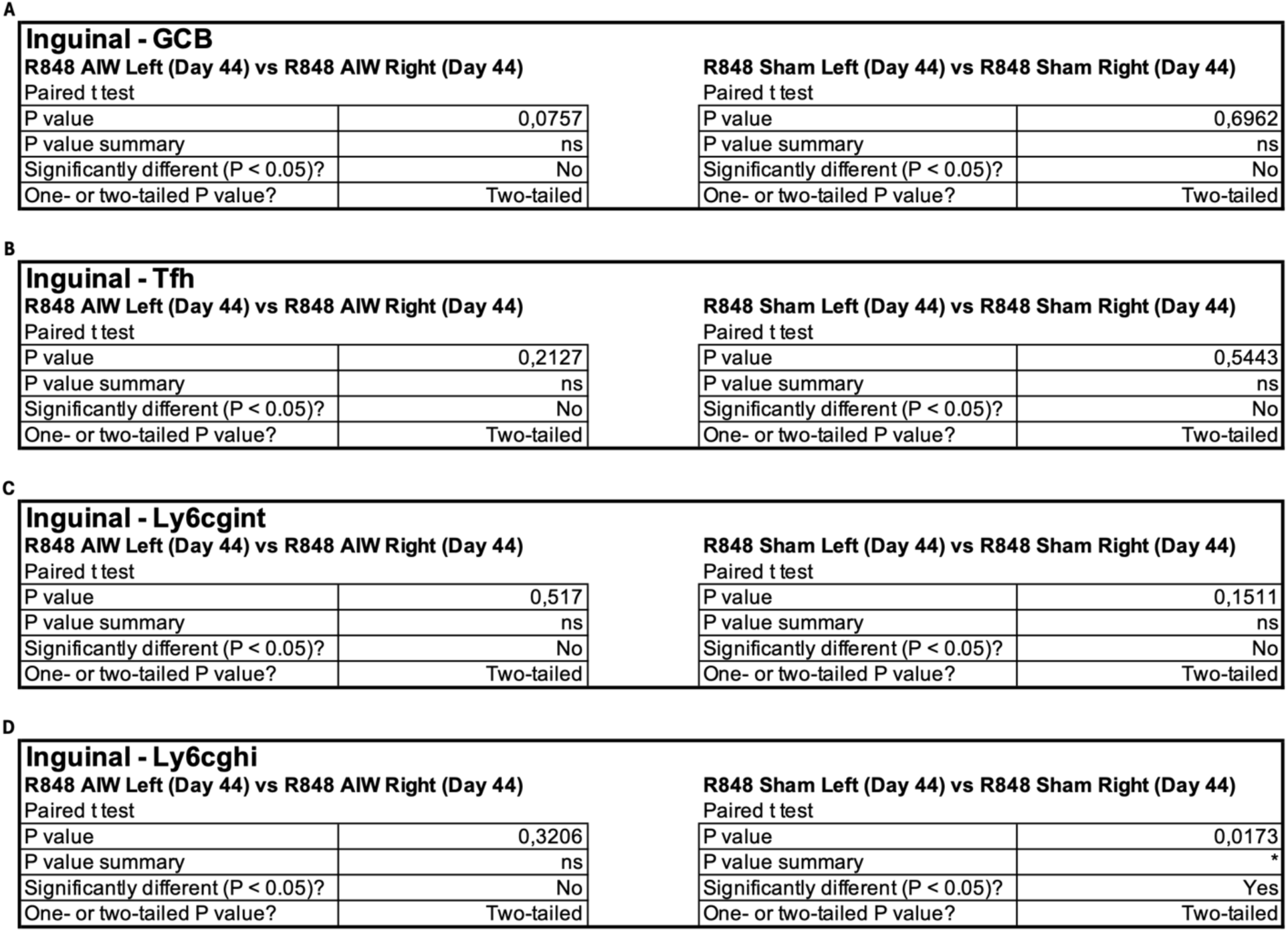
Statistical Analysis of Flow cytometry of inguinal lymph nodes of unoperated mice, Sham and AIW-operated mice to detect side differences. (A) GC B cells. (B) Tfh cells. C Ly6cg^int^ cells. D Ly6cg^hi^ cells. Test details can be found in the table.

| **Inguinal - GCB** |  |  |  |
| --- | --- | --- | --- |
| **R848 AIW Left (Day 44) vs R848 AIW Right (Day 44)** |  | **R848 Sham Left (Day 44) vs R848 Sham Right (Day 44)** |  |
| Paired t test |  | Paired t test |  |
| P value | 0,0757 | P value | 0,6962 |
| P value summary | ns | P value summary | ns |
| Significantly different (P < 0.05)? | No | Significantly different (P < 0.05)? | No |
| One- or two-tailed P value? | Two-tailed | One- or two-tailed P value? | Two-tailed |

B
| **Inguinal - Tfh** |  |  |  |
| --- | --- | --- | --- |
| **R848 AIW Left (Day 44) vs R848 AIW Right (Day 44)** |  | **R848 Sham Left (Day 44) vs R848 Sham Right (Day 44)** |  |
| Paired t test |  | Paired t test |  |
| P value | 0,2127 | P value | 0,5443 |
| P value summary | ns | P value summary | ns |
| Significantly different (P < 0.05)? | No | Significantly different (P < 0.05)? | No |
| One- or two-tailed P value? | Two-tailed | One- or two-tailed P value? | Two-tailed |

C
| **Inguinal - Ly6cgint** |  |  |  |
| --- | --- | --- | --- |
| **R848 AIW Left (Day 44) vs R848 AIW Right (Day 44)** |  | **R848 Sham Left (Day 44) vs R848 Sham Right (Day 44)** |  |
| Paired t test |  | Paired t test |  |
| P value | 0,517 | P value | 0,1511 |
| P value summary | ns | P value summary | ns |
| Significantly different (P < 0.05)? | No | Significantly different (P < 0.05)? | No |
| One- or two-tailed P value? | Two-tailed | One- or two-tailed P value? | Two-tailed |

| **Inguinal - Ly6cg<sup>hi</sup>** |  |  |  |
| --- | --- | --- | --- |
| **R848 AIW Left (Day 44) vs R848 AIW Right (Day 44)** |  | **R848 Sham Left (Day 44) vs R848 Sham Right (Day 44)** |  |
| Paired t test |  | Paired t test |  |
| P value | 0,3206 | P value | 0,0173 |
| P value summary | ns | P value summary | * |
| Significantly different (P < 0.05)? | No | Significantly different (P < 0.05)? | Yes |
| One- or two-tailed P value? | Two-tailed | One- or two-tailed P value? | Two-tailed |

**Supplementary Table 2.** Antibodies used for Flow Cytometry.

| Antibody | Dilution | Vendor | Reference Number |
| --- | --- | --- | --- |
| B220-BV510 | 1/500 | BD Horizon | 563103 |
| CD4-Qdot605 | 1/500 | Thermo Fisher Scientific | Q10092 |
| CD8-PerCP-Cy5.5 | 1/500 | BD Pharmingen | 565310 |
| CD38-PE-Cy7 | 1/500 | Nordic Biosite | 102718 |
| CD95-PE | 1/200 | BD Pharmingen | 554258 |
| CD138-BV650 | 1/500 | BD Horizon | 564068 |
| CXCR5-Biotin | 1/200 | BD Biosciences | 551960 |
| Streptavidin-BV421 | 1/200 | BD Biosciences | 563259 |
| CD25-BV786 | 1/200 | BD Biosciences | 564023 |
| Viability Dye eFluor780 | 1/2000 | Invitrogen™ eBioscience™ | 65-0865-18 |
| Ly6g/c-APC-R700 | 1/500 | BD Pharmingen | 565510 |
| GL7-AlexaFluo647 | 1/500 | BD Pharmingen | 561529 |

## References

1. S. M. Lewis, A. Williams, S. C. Eisenbarth, Structure and function of the immune system in the spleen. Sci Immunol 4, (2019).

2. V. Bronte, M. J. Pittet, The spleen in local and systemic regulation of immunity. Immunity 39, 806–818 (2013).

3. T. I. Arnon, R. M. Horton, I. L. Grigorova, J. G. Cyster, Visualization of splenic marginal zone B-cell shuttling and follicular B-cell egress. Nature 493, 684–688 (2013).

4. A. Chauveau et al., Visualization of T Cell Migration in the Spleen Reveals a Network of Perivascular Pathways that Guide Entry into T Zones. Immunity 52, 794–807 e797 (2020).

5. H. S. Jeong et al., Investigation of the Lack of Angiogenesis in the Formation of Lymph Node Metastases. J Natl Cancer Inst 107, (2015).

6. E. F. J. Meijer et al., Murine chronic lymph node window for longitudinal intravital lymph node imaging. Nat Protoc 12, 1513–1520 (2017).

7. E. R. Pereira et al., Lymph node metastases can invade local blood vessels, exit the node, and colonize distant organs in mice. Science 359, 1403–1407 (2018).

8. D. J. Firl, S. E. Degn, T. Padera, M. C. Carroll, Capturing change in clonal composition amongst single mouse germinal centers. Elife 7, (2018).

9. J. T. Jacobsen et al., Expression of Foxp3 by T follicular helper cells in end-stage germinal centers. Science 373, (2021).

10. L. Ritsma et al., Surgical implantation of an abdominal imaging window for intravital microscopy. Nat Protoc 8, 583–594 (2013).

11. L. Ritsma et al., Intravital microscopy through an abdominal imaging window reveals a pre-micrometastasis stage during liver metastasis. Sci Transl Med 4, 158ra145 (2012).

12. L. Bordoni et al., Longitudinal tracking of acute kidney injury reveals injury propagation along the nephron. Nat Commun 14, 4407 (2023).

13. K. Zhang et al., In vivo two-photon microscopy reveals the contribution of Sox9+ cell to kidney regeneration in a mouse model with extracellular vesicle treatment. Journal of Biological Chemistry 295, 12203–12213 (2020).

14. I. Park, P. Kim, Stabilized Longitudinal In Vivo Cellular-Level Visualization of the Pancreas in a Murine Model with a Pancreatic Intravital Imaging Window. J Vis Exp, e62538 (2021).

15. D. L. Marvin, P. ten Dijke, L. Ritsma, An Experimental Liver Metastasis Mouse Model Suitable for Short and Long-Term Intravital Imaging. Current Protocols 1, (2021).

16. D. E. DeTemple et al., Longitudinal imaging and femtosecond laser manipulation of the liver: How to generate and trace single-cell-resolved micro-damage in vivo. PLoS One 15, e0240405 (2020).

17. J. Herz, B. H. Zinselmeyer, D. B. McGavern, Two-Photon Imaging of Microbial Immunity in Living Tissues. Microscopy and Microanalysis 18, 730–741 (2012).

18. M. Yokogawa et al., Epicutaneous application of toll-like receptor 7 agonists leads to systemic autoimmunity in wild-type mice: a new model of systemic Lupus erythematosus. Arthritis Rheumatol 66, 694–706 (2014).

19. R. Berland et al., Toll-like receptor 7-dependent loss of B cell tolerance in pathogenic autoantibody knockin mice. Immunity 25, 429–440 (2006).

20. S. R. Christensen et al., Toll-like receptor 7 and TLR9 dictate autoantibody specificity and have opposing inflammatory and regulatory roles in a murine model of lupus. Immunity 25, 417–428 (2006).

21. R. A. Herlands, S. R. Christensen, R. A. Sweet, U. Hershberg, M. J. Shlomchik, T cell-independent and toll-like receptor-dependent antigen-driven activation of autoreactive B cells. Immunity 29, 249–260 (2008).

22. C. M. Lau et al., RNA-associated autoantigens activate B cells by combined B cell antigen receptor/Toll-like receptor 7 engagement. J Exp Med 202, 1171–1177 (2005).

23. G. J. Brown et al., TLR7 gain-of-function genetic variation causes human lupus. Nature, (2022).

24. R. A. Elsner, M. J. Shlomchik, Germinal Center and Extrafollicular B Cell Responses in Vaccination, Immunity, and Autoimmunity. Immunity 53, 1136–1150 (2020).

25. S. A. Jenks, K. S. Cashman, M. C. Woodruff, F. E. Lee, I. Sanz, Extrafollicular responses in humans and SLE. Immunol Rev 288, 136–148 (2019).

26. L. F. Voss et al., The extrafollicular response is sufficient to drive initiation of autoimmunity and early disease hallmarks of lupus. Front Immunol 13, 1021370 (2022).

27. S. E. Degn et al., Clonal Evolution of Autoreactive Germinal Centers. Cell 170, 913–926 e919 (2017).

28. I. G. Luzina et al., Spontaneous formation of germinal centers in autoimmune mice. J Leukoc Biol 70, 578–584 (2001).

29. P. P. Domeier, S. L. Schell, Z. S. Rahman, Spontaneous germinal centers and autoimmunity. Autoimmunity 50, 4–18 (2017).

30. A. Rahman, D. A. Isenberg, Systemic lupus erythematosus. N Engl J Med 358, 929–939 (2008).

31. C. Soni et al., B cell-intrinsic TLR7 signaling is essential for the development of spontaneous germinal centers. J Immunol 193, 4400–4414 (2014).

32. K. Green et al., B Cell Intrinsic STING Signaling Is Not Required for Autoreactive Germinal Center Participation. Front Immunol 12, 782558 (2021).

33. S. W. Jackson et al., Opposing impact of B cell-intrinsic TLR7 and TLR9 signals on autoantibody repertoire and systemic inflammation. J Immunol 192, 4525–4532 (2014).

34. A. Das et al., Follicular Dendritic Cell Activation by TLR Ligands Promotes Autoreactive B Cell Responses. Immunity 46, 106–119 (2017).

35. P. P. Domeier et al., B-Cell-Intrinsic Type 1 Interferon Signaling Is Crucial for Loss of Tolerance and the Development of Autoreactive B Cells. Cell Rep 24, 406–418 (2018).

36. D. L. Thibault et al., Type I interferon receptor controls B-cell expression of nucleic acid-sensing Toll-like receptors and autoantibody production in a murine model of lupus. Arthritis Res Ther 11, R112 (2009).

37. G. D. Victora, M. C. Nussenzweig, Germinal Centers. Annu Rev Immunol, (2022).

38. L. T. Krimpenfort, S. E. Degn, B. A. Heesters, The follicular dendritic cell: At the germinal center of autoimmunity? Cell Rep 43, 113869 (2024).

39. I. M. Schiessl, K. Fremter, J. L. Burford, H. Castrop, J. Peti-Peterdi, Long-Term Cell Fate Tracking of Individual Renal Cells Using Serial Intravital Microscopy. Methods Mol Biol 2150, 25–44 (2020).

40. D. Sardella et al., Serial intravital 2-photon microscopy and analysis of the kidney using upright microscopes. Frontiers in Physiology 14, (2023).

41. L. Ritsma et al., Surgical implantation of an abdominal imaging window for intravital microscopy. Nature Protocols 8, 583–594 (2013).

42. L. Ritsma et al., Intravital Microscopy Through an Abdominal Imaging Window Reveals a Pre-Micrometastasis Stage During Liver Metastasis. Science Translational Medicine 4, 158ra145–158ra145 (2012).

43. J. M. Anderson, A. Rodriguez, D. T. Chang, Foreign body reaction to biomaterials. Seminars in Immunology 20, 86–100 (2008).

44. R. Konradi, C. Acikgoz, M. Textor, Polyoxazolines for Nonfouling Surface Coatings — A Direct Comparison to the Gold Standard PEG. Macromolecular Rapid Communications 33, 1663–1676 (2012).

45. Y. Chen et al., Comparative assessment of the stability of nonfouling poly(2-methyl-2-oxazoline) and poly(ethylene glycol) surface films: An in vitro cell culture study. Biointerphases 9, 031003 (2014).

46. Â. Serrano, S. Zürcher, S. Tosatti, N. D. Spencer, Imparting Nonfouling Properties to Chemically Distinct Surfaces with a Single Adsorbing Polymer: A Multimodal Binding Approach. Macromolecular Rapid Communications 37, 622–629 (2016).

47. S. R. Meyers, M. W. Grinstaff, Biocompatible and Bioactive Surface Modifications for Prolonged In Vivo Efficacy. Chemical Reviews 112, 1615–1632 (2012).

48. M. Yokogawa et al., Epicutaneous Application of Toll-like Receptor 7 Agonists Leads to Systemic Autoimmunity in Wild-Type Mice: A New Model of Systemic Lupus Erythematosus. Arthritis & Rheumatology 66, 694–706 (2014).

49. J. P. Smith, G. F. Burton, J. G. Tew, A. K. Szakal, Tinigible Body Macrophages in Regulation of Germinal Center Reactions. Journal of Immunology Research 6, 038923 (1998).

50. Y. Chung et al., Follicular regulatory T cells expressing Foxp3 and Bcl-6 suppress germinal center reactions. Nature Medicine 17, 983–988 (2011).

51. F. Helmchen, W. Denk, Deep tissue two-photon microscopy. Nature Methods 2, 932–940 (2005).

52. T. G. Phan, A. Bullen, Practical intravital two-photon microscopy for immunological research: faster, brighter, deeper. Immunology & Cell Biology 88, 438–444 (2010).

53. P. Luu, S. E. Fraser, F. Schneider, More than double the fun with two-photon excitation microscopy. Communications Biology 7, 364 (2024).

54. J. Schindelin et al., Fiji: an open-source platform for biological-image analysis. Nature Methods 9, 676–682 (2012).

55. J. Chapman, A. Goyal, A. M. Azevedo, in StatPearls [Internet]. (StatPearls Publishing, 2021).

56. L. F. Voss et al., The extrafollicular response is sufficient to drive initiation of autoimmunity and early disease hallmarks of lupus. Frontiers in Immunology 13, 1021370–1021370 (2022).

57. A. K. Grootveld et al., Apoptotic cell fragments locally activate tingible body macrophages in the germinal center. Cell 186, 1144–1161.e1118 (2023).

58. A. Chauveau et al., Visualization of T Cell Migration in the Spleen Reveals a Network of Perivascular Pathways that Guide Entry into T Zones. Immunity 52, 794–807.e797 (2020).

59. D. Liu et al., Dynamic encounters with red blood cells trigger splenic marginal zone B cell retention and function. Nature Immunology 25, 142–154 (2024).

60. T. Hanawa, Biocompatibility of titanium from the viewpoint of its surface. Science and Technology of Advanced Materials 23, 457–472 (2022).

61. T. Poole, Happy animals make good science. Laboratory Animals 31, 116–124 (1997).

62. M. J. Prescott, K. Lidster, Improving quality of science through better animal welfare: the NC3Rs strategy. Lab Animal 46, 152–156 (2017).

63. K. Viswanathan, F. S. Dhabhar, Stress-induced enhancement of leukocyte trafficking into sites of surgery or immune activation. Proceedings of the National Academy of Sciences 102, 5808–5813 (2005).

64. M. Alieva, L. Ritsma, R. J. Giedt, R. Weissleder, J. van Rheenen, Imaging windows for long-term intravital imaging. IntraVital 3, e29917 (2014).

65. M. P. S. Dunphy, D. Entenberg, R. Toledo-Crow, S. M. Larson, In vivo microcartography and subcellular imaging of tumor angiogenesis: A novel platform for translational angiogenesis research. Microvascular Research 78, 51–56 (2009).

66. D. U. Pizzagalli et al., Leukocyte Tracking Database, a collection of immune cell tracks from intravital 2-photon microscopy videos. Scientific Data 5, 180129 (2018).

67. D. Soulet, J. Lamontagne-Proulx, B. AubÉ, D. Davalos, Multiphoton intravital microscopy in small animals: motion artefact challenges and technical solutions. Journal of Microscopy 278, 3–17 (2020).

68. M. R. Arbuckle et al., Development of Autoantibodies before the Clinical Onset of Systemic Lupus Erythematosus. New England Journal of Medicine 349, 1526–1533 (2003).

69. T. Shay et al., Conservation and divergence in the transcriptional programs of the human and mouse immune systems. Proceedings of the National Academy of Sciences 110, 2946–2951 (2013).

70. J. Seok et al., Genomic responses in mouse models poorly mimic human inflammatory diseases. Proceedings of the National Academy of Sciences 110, 3507–3512 (2013).

71. S. M. Lewis, A. Williams, S. C. Eisenbarth, Structure and function of the immune system in the spleen. Sci Immunol 4, eaau6085 (2019).

72. B. S. Steiniger, Human spleen microanatomy: why mice do not suffice. Immunology 145, 334–346 (2015).

73. J. M. Kim, J. P. Rasmussen, A. Y. Rudensky, Regulatory T cells prevent catastrophic autoimmunity throughout the lifespan of mice. Nature Immunology 8, 191–197 (2007).

74. G. D. Victora et al., Germinal Center Dynamics Revealed by Multiphoton Microscopy with a Photoactivatable Fluorescent Reporter. Cell 143, 592–605 (2010).

75. R. Ogaki et al., Temperature-Induced Ultradense PEG Polyelectrolyte Surface Grafting Provides Effective Long-Term Bioresistance against Mammalian Cells, Serum, and Whole Blood. Biomacromolecules 13, 3668–3677 (2012).

76. S. Ali, S. Malthe von Tangen, L. Sara Hvidbjerg, S. S. Duncan, Microplate Format Protein Nanopatterning for High-Throughput Screening of Cellular Microenvironments. bioRxiv, 2023.2011.2019.567703 (2023).

77. S. Carsen, P. Marius, Cellpose3: one-click image restoration for improved cellular segmentation. bioRxiv, 2024.2002.2010.579780 (2024).

78. G. Cox et al., 3-Dimensional imaging of collagen using second harmonic generation. Journal of Structural Biology 141, 53–62 (2003).

79. R. Zipfel Warren et al., Live tissue intrinsic emission microscopy using multiphoton-excited native fluorescence and second harmonic generation. Proceedings of the National Academy of Sciences 100, 7075–7080 (2003).

80. N. Chiaruttini, O. Burri, Multi Stack Montage. https://github.com/BIOP/ijp-multi-stack-montage.

81. L. Ritsma, N. Vrisekoop, J. van Rheenen, In vivo imaging and histochemistry are combined in the cryosection labelling and intravital microscopy technique. Nature Communications 4, 2366 (2013).

82. J. A. Bogovic, P. Hanslovsky, A. Wong, S. Saalfeld, in 2016 IEEE 13th international symposium on biomedical imaging (ISBI). (IEEE, 2016), pp. 1123–1126.

